# The role of the glutamate-glutamine cycle in synaptic transmission during ischemia and recovery

**DOI:** 10.1101/2025.11.10.687543

**Authors:** Hannah van Susteren, Christine R. Rose, Michel J.A.M. van Putten, Hil G.E. Meijer

## Abstract

Cerebral ischemia impairs neuronal and glial function, ranging from synaptic failure to irreversible damage. The effects of ischemia on excitatory synaptic transmission remain incompletely understood. Here, we present a detailed biophysical model including the first full implementation of the glutamate-glutamine cycle (GG-cycle), which is essential for proper functioning of glutamatergic synapses.

We simulate a presynaptic neuron and an astrocyte in a finite extracellular space (ECS), surrounded by an oxygen bath, as a proxy for energy supply. The model includes ionic currents with corresponding channels and transporters such as the sodium-potassium ATPase. To model synaptic transmission, we combine calcium-dependent glutamate release, its uptake by the sodium-dependent excitatory amino acid transporters (EAATs), and the GG-cycle, including glutamine synthesis.

We simulate ischemia by blocking energy supply completely. The neuron enters a depolarization block, ion concentrations reach pathological values, and glutamate accumulates in the ECS while glutamate release is disrupted. Surprisingly, we found that synaptic transmission failure was not primarily caused by excessive glutamate release nor by failure of glutamine synthetase. Instead, it mainly resulted from EAAT dysfunction, driven by the collapse of the sodium gradient. Enhancing glutamate clearance alone was insufficient for recovery of synaptic transmission. However, inhibition of the voltage-gated Na^+^ channels restored ion gradients, recovered glutamate uptake, and re-enabled glutamate release.

Taken together, our study highlights the critical role of ion homeostasis, in particular the sodium gradient, in maintaining synaptic function during metabolic stress. Moreover, the model provides a better understanding of synaptic transmission failure and potential recovery strategies during ischemia.

## 1 Introduction

Synaptic transmission is essential for normal brain function and depends critically on continuous energy supply. During cerebral ischemia, synaptic transmission is one of the first processes to fail (Hofmeijer & van Putten, 2012), affecting both excitatory and inhibitory neurotransmission. This disruption includes dysregulation of glutamate homeostasis, which may lead to excitotoxicity, and ultimately neuronal injury or death (Choi, 1988; Martinez Cruz et al., 2017; Olloquequi et al., 2018).

Under physiological conditions, excess extracellular glutamate is primarily cleared by the astrocytic excitatory amino acid transporters (EAATs) (Mahmoud et al., 2019). The main pathway for astrocytic glutamate is the glutamate-glutamine cycle (GG-cycle) (Bak et al., 2006; Lemberg & Alejandra Fernández, 2009; Waagepetersen et al., 2005) in which astrocytic glutamate is converted into glutamine by the energy-dependent enzyme glutamine synthetase (Bak et al., 2006; Lieth et al., 2001; Petito et al., 1992). Following synthesis, glutamine is transported out of astrocytes and taken up by neurons (S. Bröer & Brookes, 2001; Chaudhry et al., 2002; Mackenzie et al., 2003; Todd et al., 2017; Yao et al., 2000), where it is converted back into glutamate (Kvamme et al., 2000, 2001). In the neuron, vesicle recycling replenishes the neuronal depot, enabling glutamate release during neuronal activity. While the essential role of the glutamate-glutamine cycle in regulating synaptic transmission under physiological conditions is well understood, its role in disrupted synaptic transmission during ischemia and subsequent recovery remains unclear.

Biophysical models are powerful tools for exploring the complex interactions underlying synaptic transmission failure during ischemia, as they enable simultaneous analysis of neurotransmitter dynamics and associated ion fluxes. We build on previous computational studies of single synapses focusing on glutamate dynamics. For instance, studies on single synapses have demonstrated the importance of astrocytic glutamate uptake and its impact on postsynaptic receptors (Allam et al., 2012; Dronne et al., 2007). Other studies have explored the cycling rates of glutamate and glutamine in isolated systems (Shen, 2013).

Subsequent research has extended these models to include ion dynamics in the tripartite synapse, e.g., highlighting the critical consequences of increased astrocytic sodium concentrations due to increased EAAT activity, which can reverse the action of the sodium-calcium exchanger (NCX) (Breslin et al., 2018). Additionally, impaired glutamate uptake has been shown to prolong the duration of spreading depression and prevent recovery (Hübel et al., 2017), while elevated astrocytic glutamate concentrations alter the temporal profile of glutamate in the extracellular space (Flanagan et al., 2018). Finally, detailed modeling of the tripartite synapse including ion and glutamate dynamics during ischemia identified key factors for reversible or irreversible synaptic damage (Kalia et al., 2021).

Although these studies provide valuable insights on synaptic transmission, none have incorporated the full dynamics of the GG-cycle alongside ion dynamics. For a more comprehensive understanding of synaptic failure during metabolic stress, we now integrated this essential component. In this work, we present a detailed biophysical model that includes the full dynamics of the glutamate-glutamine cycle, offering the first complete simulation of both glutamate dynamics and ion fluxes in three synaptic compartments.

Our model is comprised of a neuron, an astrocyte, and a finite extracellular space (ECS). Ion movement is based on the principle of electrodiffusion. Synaptic transmission includes calcium-dependent vesicle recycling, and a novel implementation of the GG-cycle. Cell swelling is modelled using passive water influx driven by osmotic pressure gradients. Finally, to simulate varying degrees of energy deprivation, we incorporate oxygen dynamics along with an external oxygen bath that allows diffusion into the ECS. With this model, we examine the regulation of synaptic transmission by the glutamate–glutamine cycle during ischemia and recovery.

## 2 Results

Before exploring ischemic conditions, we demonstrate the importance of the glutamate-glutamine cycle for synaptic transmission in physiological conditions. This is followed by simulations of moderate and severe ischemia, demonstrating the failure of synaptic transmission and showing glutamate accumulation. Subsequently, we examine the mechanisms contributing to glutamate accumulation. Lastly, we evaluate potential strategies to restore synaptic transmission.

### 2.1 Glutamate dynamics in physiology and during ischemia

#### 2.1.1 Glutamate dynamics during physiological conditions

Figure 1 shows glutamate and glutamine concentrations during rest and external electrical stimulation used to generate a burst of action potentials. In response to stimulation, the neuron depolarizes, calcium enters the cell, and glutamate is released from the neuronal depot into the ECS, showing glutamate peaks as expected. By blocking glutamine synthetase, we simulate the effect of methionine sulfoximine (MSO) (Rothstein & Tabakoff, 1984). This block results in the depletion of all glutamine concentrations as there is no glutamine supply through the GG-cycle, while astrocytic and neuronal glutamate accumulate due to the block. Stimulating the neuron under these conditions results in a reduction in glutamate release (black trace in Figure 1).

**Figure 1.**
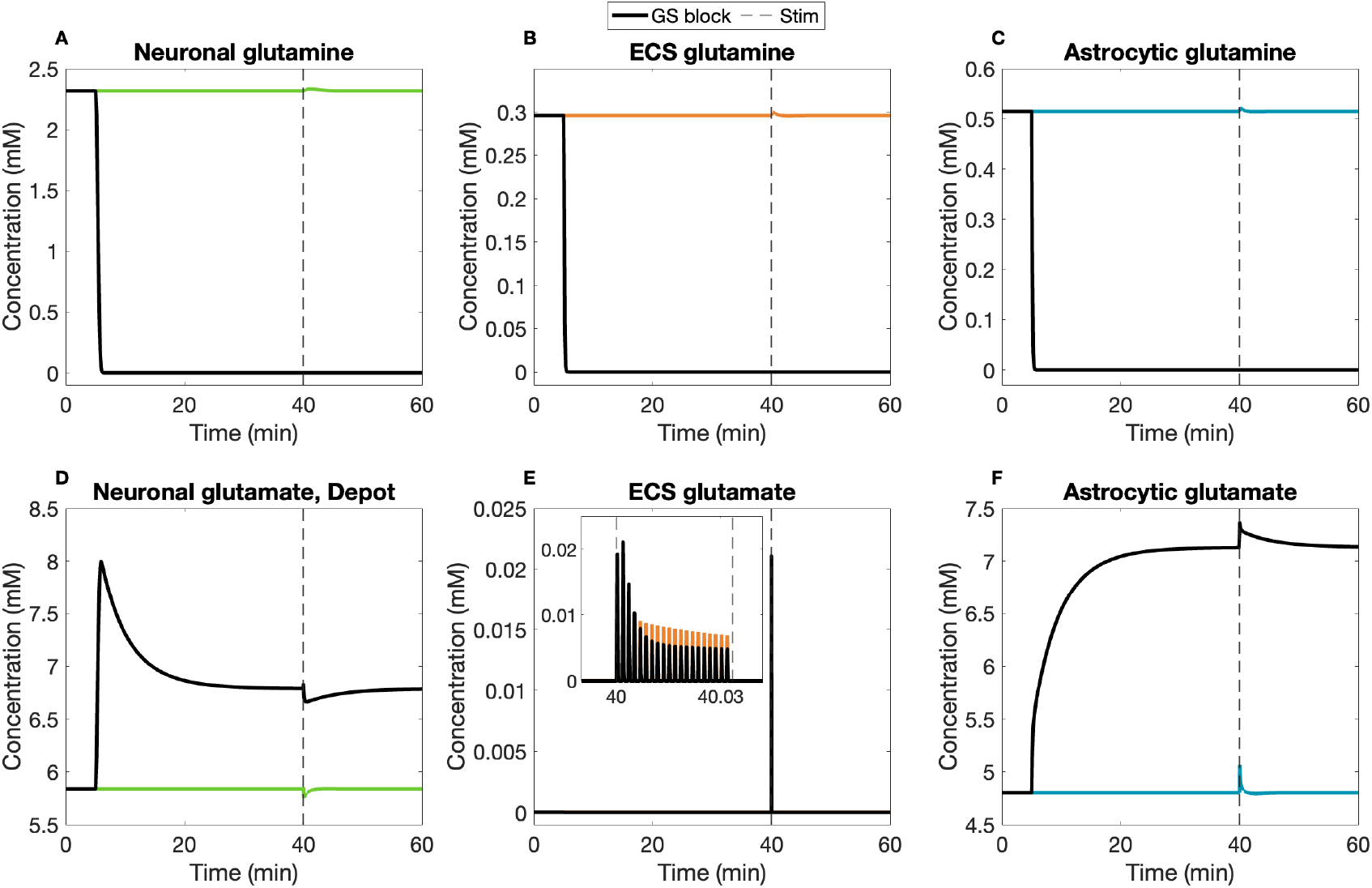
Glutamate and glutamine concentrations in physiological conditions with glutamate synthetase inhibition. The glutamine (A,B,C) and glutamate (D,E,F) concentrations in the neuron (green), ECS (orange) and astrocyte (blue) are shown in physiological conditions. Blockade of glutamine synthetase at t=5 minutes (black lines) leads to depletion of glutamine concentrations, an increased neuronal glutamate depot and increased astrocytic glutamate. External stimulation at t=40 min during 2 seconds results in a burst of action potentials, with release of glutamate gradually decaying in amplitude. (E) Glutamate release is decreased during GS inhibition compared to physiological conditions.

#### 2.1.2 Glutamate Dynamics During Moderate Ischemia

To simulate different ischemic conditions, we vary the oxygen concentration in the bath. For moderate ischemia, we lower the oxygen concentration in the bath to 75% for 55 minutes, see Figure 2. During 55 minutes of moderate ischemia, the oxygen concentration in the ECS declines due to restricted oxygen availability from the bath and energy consumption by neurons and astrocytes. The sodium-potassium ATPase (NKA) activity decreases, leading to temporary changes in ion concentrations. This results in an increase in the membrane potentials and simultaneously an increase in extracellular glutamate due to decreased EAAT activity. When oxygen concentration in the bath is restored, membrane potentials and extracellular glutamate all return to their pre-ischemia levels. To verify physiological dynamics after moderate ischemia, the neuron is stimulated with an electrical current pulse at t=80 min for 2 ms, which is simulated using a square-wave sodium current. Action potentials are generated, and the calcium-dependent fusion of synaptic vesicles is induced, resulting in glutamate release from the neuron into the ECS. This indicates that physiological dynamics are restored and that the system can recover from moderate ischemia.

**Figure 2.**
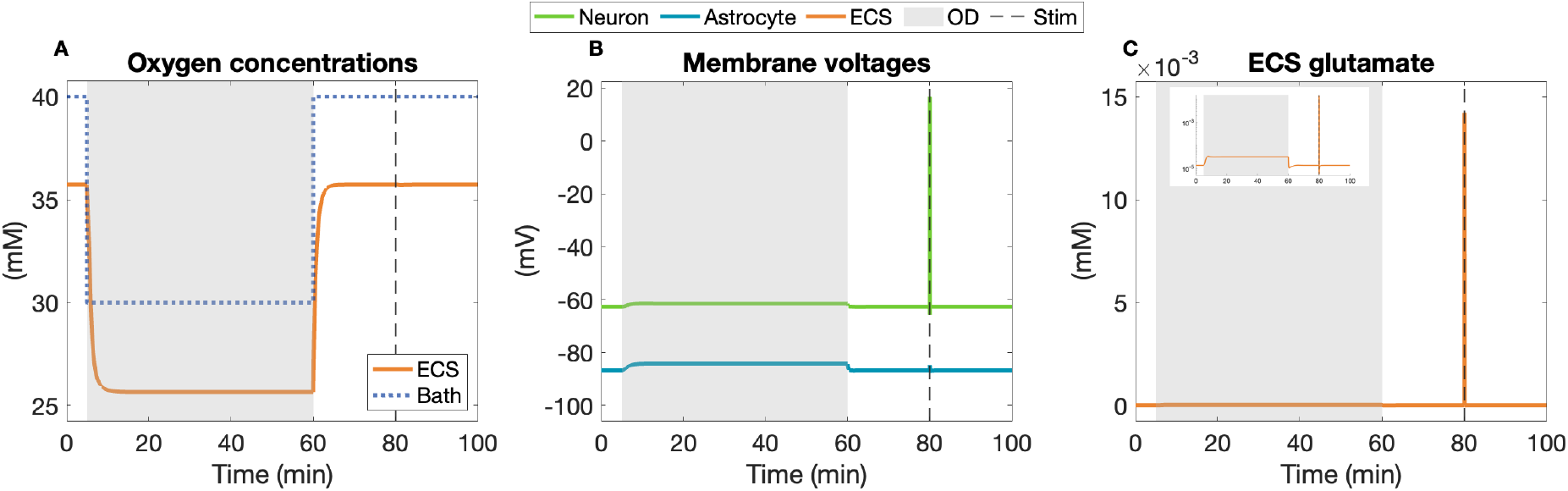
Moderate ischemia followed by brief electrical stimulation. (A) The oxygen concentration in the bath is reduced to 75% of baseline between t=5 and t=60 minutes. The extracellular oxygen concentration decreases during moderate ischemia and returns to baseline afterwards. (B) Membrane potentials increase marginally. Action potentials are generated in response to an electrical current pulse at t=80 minutes. (C) The extracellular glutamate concentration remains low during moderate ischemia. Glutamate is released in response to the action potential.

#### 2.1.3 Glutamate Dynamics During Severe Ischemia

We simulate severe ischemia by reducing the oxygen level in the bath to zero for 55 minutes, shown in Figure 3. In response to complete energy depletion, the extracellular oxygen concentration transiently drops to zero (Figure 3A), leading to reduced activity of all energy-dependent processes, including the neuronal and astrocytic NKA. Without functioning NKA, the neuron depolarizes, transits to a period of spontaneous action potential firing, and ultimately reaches a pathological depolarized state. Concurrently, intracellular of sodium and calcium and extracellular potassium increase (Figure 3D,E,F). These ion imbalances lead to build-up in osmotic pressure, causing water influx and subsequent swelling of both the neuron and astrocyte (Figure 3C).

**Figure 3.**
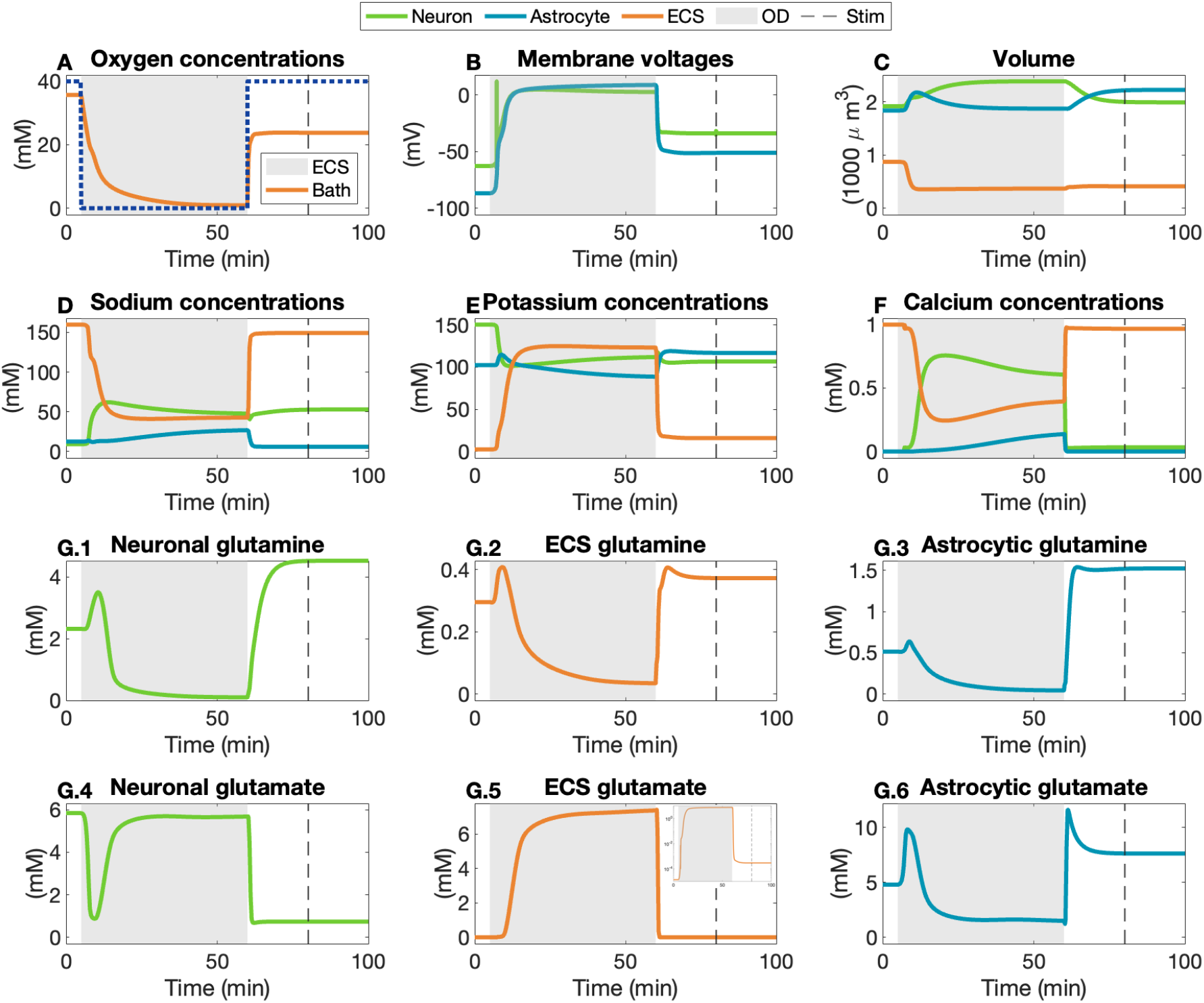
Severe ischemia followed by electrical stimulation. (A) Oxygen in the bath is completely depleted between t=5 and t=60 minutes. In response, extracellular oxygen depletes. (B) The neuron enters a depolarization block, and settles in a pathological state. (C) The neuronal and astrocytic volumes increase. (D) Neuronal and astrocytic sodium increase. (E) Extracellular potassium increases significantly. (F) Neuronal and astrocytic calcium increase. (G) Dynamics of the full glutamate-glutamine cycle during ischemia. (G.1-3) show glutamine concentrations in the neuron, ECS and astrocyte, respectively. (G.4-6) show glutamate concentrations in the neuron, ECS and astrocyte, respectively. (G.5) Glutamate accumulation in response to five minutes of severe ischemia. Severe ischemia results in disturbed ion homeostasis and synaptic transmission failure. Upon stimulation at t=80 minutes, no action potential is generated and glutamate release is absent.

Figure 3G shows the effects of severe ischemia on the dynamics of the glutamate-glutamine cycle. Most noteworthy, extracellular glutamate increases drastically up to 7.5 mM (Figure 3G.5). In the first minutes of ischemia, a fraction of extracellular glutamate is taken up by the astrocytic EAAT, which exhibits a rapid and transient peak at the start of ischemia, see Figure 4A. Initially, glutamate is still converted into glutamine in the astrocyte and transported to the ECS and neuron, showing transient increases in glutamine concentrations (Figure 3G.1-3). After these transients, both the EAAT and glutamine synthetase cease activity, see Figure 4B. As glutamate uptake failure precedes glutamine synthetase malfunction, this results in depletion of astrocytic glutamate and subsequently also astrocytic glutamine. The other components of the GG-cycle remain functional, eventually leading to depletion of all glutamine concentrations. When glutamate release from the neuronal stores ceases and glutamine continues to be transported into the neuron for conversion, the neuronal glutamate concentration increases.

**Figure 4.**
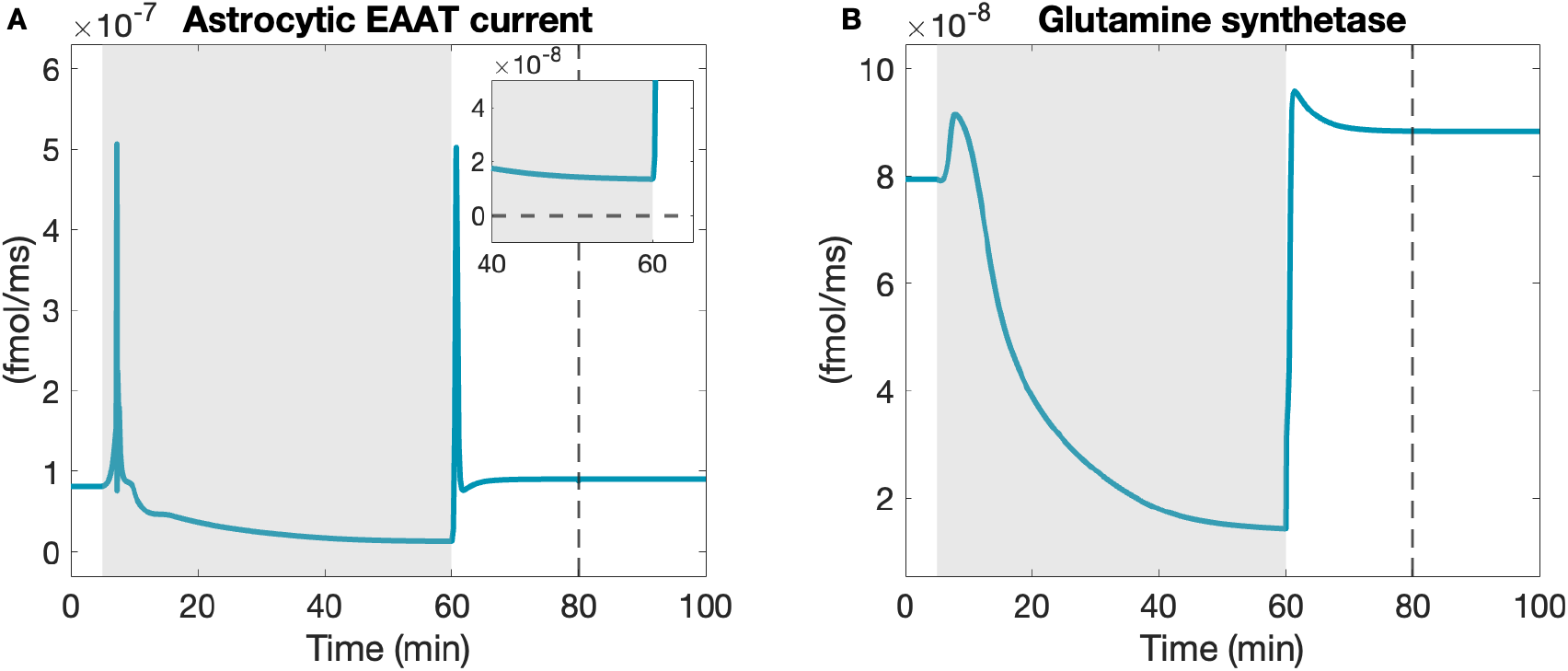
The astrocytic EAAT current (A) and glutamine synthetase current (B) during ischemia. Oxygen deprivation occurs between t=5 to t=60 minutes, with neuronal stimulation at t=80 minutes. Failure of glutamate uptake precedes malfunction of glutamine synthesis. The astrocytic EAAT current exhibits a rapid increase in activity before malfunctioning, and does not reverse direction (as shown in the inset). Glutamine synthetase gradually reduces activity until malfunction.

Having observed the effects of severe ischemia, we study how the system reacts when we restore the oxygen supply to baseline at t=60 minutes. In contrast to the scenario with moderate ischemia, the extracellular oxygen concentration does not recover (Figure 3A). Neuronal and astrocytic NKA cannot clear the excess intracellular sodium; both the neuron and astrocyte remain depolarized, and cell swelling persists (Figure 3C-F). Similarly, the glutamate-glutamine cycle does not recover after energy supply has returned (Figure 3G). An initial increase in glutamate uptake leads to an increase in astrocytic glutamate. Despite limited energy available, glutamine synthetase is able to resume activity and convert glutamate into glutamine, increasing all glutamine concentrations. The glutamate supply to the depot diminishes following ischemia, and the neuronal glutamate depot gets depleted. Lastly, the neuronal and astrocytic EAAT are unable to clear the remaining glutamate from the extracellular space after ischemia, resulting in a residual concentration of 0.3 *µ*M.

To test synaptic transmission following ischemia, the neuron is stimulated with a short external current at t=80 minutes. No action potentials are generated, and glutamate release is absent. In conclusion, 55 minutes of severe ischemia result in complete synaptic transmission failure.

### 2.2 Glutamate accumulation

Our simulations show that severe ischemia results in irreversible synaptic transmission failure and glutamate accumulation. We explore three candidate mechanisms that may be involved in the accumulation of glutamate: a reduction of the ATP-dependent glutamine synthetase, increased release of glutamate, or a reduction in its clearance.

#### 2.2.1 Role of glutamine synthetase

We hypothesized that the energy dependence of glutamine synthetase may be a significant factor contributing to the glutamate build-up during ischemia (Lemberg & Alejandra Fernández, 2009; Petito et al., 1992). To test this hypothesis, we simulate an energy-independent variant of glutamate synthetase, shown in Figure 5. Glutamine synthetase remains more active when it does not depend on energy, resulting in a lower astrocytic glutamate concentration. Importantly, the astrocytic EAAT current still remains impaired. Astrocytic glutamine surplus enters the glutamate–glutamine cycle and is eventually converted back to glutamate, which accumulates in the extracellular space since glutamate uptake remains impaired. Therefore, an energy-independent variant of glutamine synthetase aggravates glutamate accumulation in the extracellular space. This demonstrates that the malfunction of ATP-dependent glutamine synthetase is not the main cause of glutamate accumulation during ischemia.

**Figure 5.**
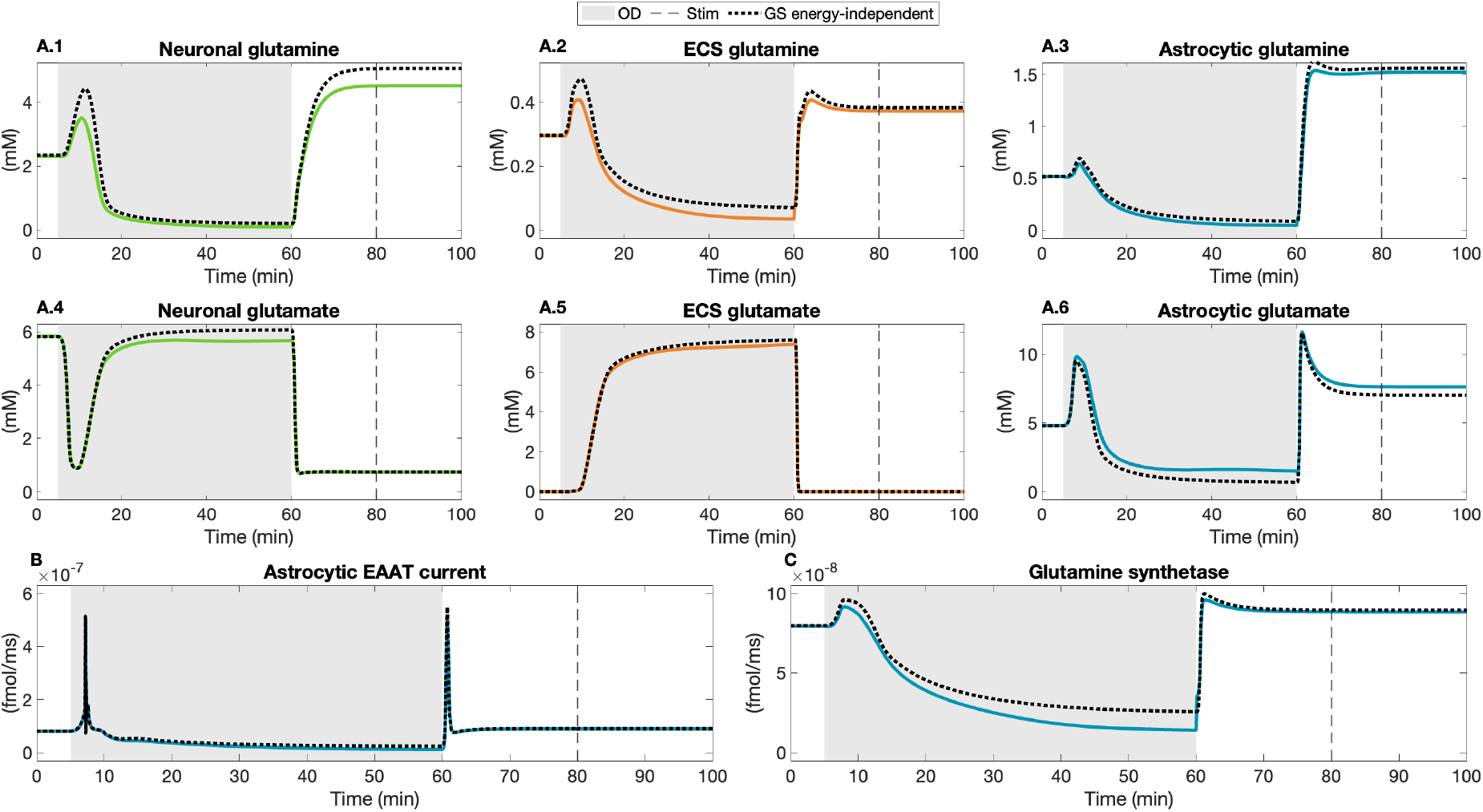
Simulation of energy-independent glutamine synthetase. Black dashed lines indicate the new simulation, colored lines indicate severe ischemia in default conditions. Energy-independent glutamine synthetase causes increase in the extracellular glutamate accumulation. (A.1-6) shows the glutamate and glutamine dynamics. (B) The astrocytic EAAT transporter current. (C) The glutamine synthetase current.

#### 2.2.2 Increased glutamate release

Glutamate accumulation could result from increased glutamate release, either through neuronal activity or EAAT reversal. A burst of neuronal action potentials is observed during the first minutes of ischemia. During the burst, glutamate is actively released through calcium-dependent exocytosis. However, the extracellular glutamate concentration only increases up to 0.01 mM during the neuronal activity (Figure 6). In the remaining period of oxygen deprivation, the glutamate concentration continues to rise to 7.5 mM, reaching extremely high levels. Moreover, no glutamate is released by the EAAT transporter since it does not reverse under ischemic conditions. Thus, neither synaptic glutamate release nor EAAT reversal contribute substantially to glutamate accumulation.

**Figure 6.**
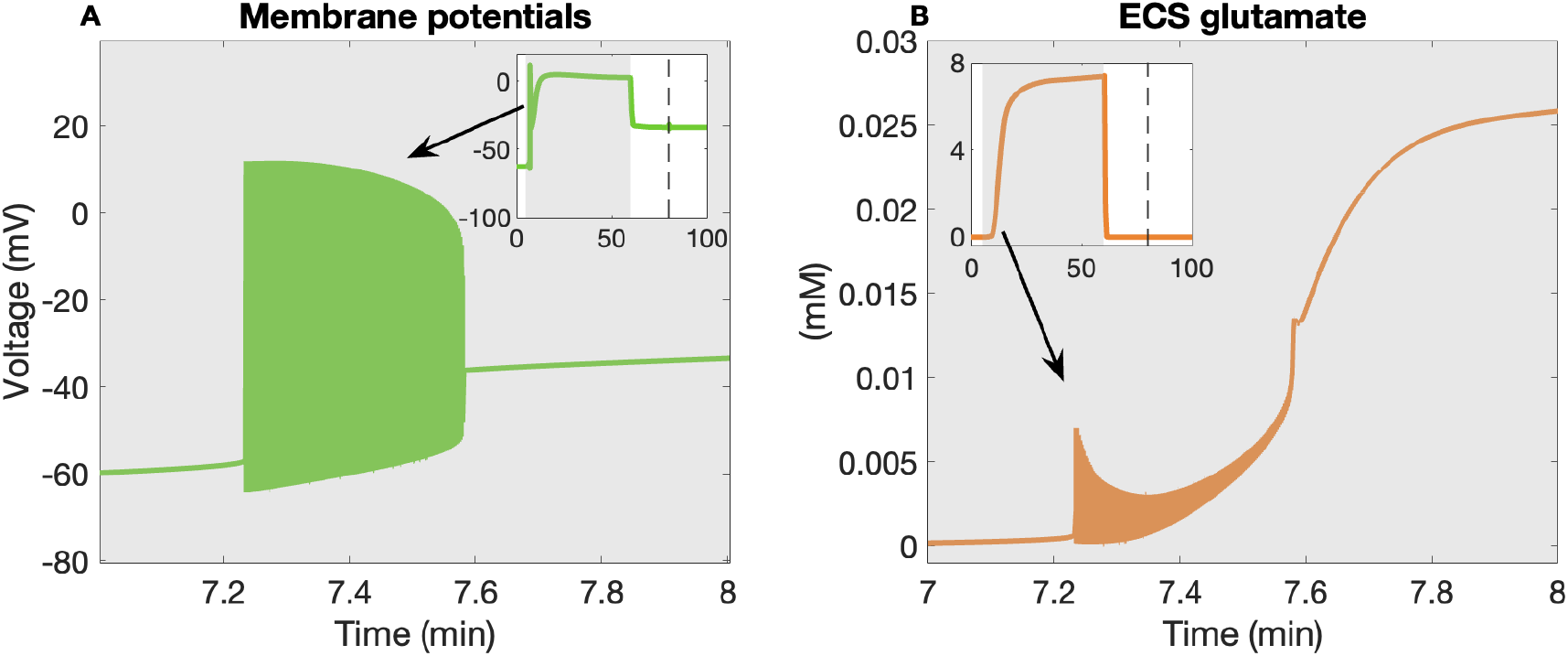
Glutamate exocytosis during ischemia. The neuronal membrane potential (A) and the extracellular glutamate concentration (B) during severe ischemia, with a detailed view of the first few minutes. The rise in extracellular glutamate during initial neuronal action potential activity is minimal.

#### 2.2.3 Role of astrocytic EAATs

Since elevated extracellular glutamate levels cannot be fully explained by increased glutamate release, EAAT reversal, or impaired glutamine synthetase activity, we hypothesize that the primary cause is impaired glutamate uptake by the astrocytic EAATs. To verify this, we simulate our ischemia scenario with the EAAT conductance being doubled or tripled, shown in Figure 7. The increased EAAT activity results in only a marginal reduction of extracellular glutamate. We infer that impairment of the EAATs plays an important role in glutamate accumulation.

**Figure 7.**
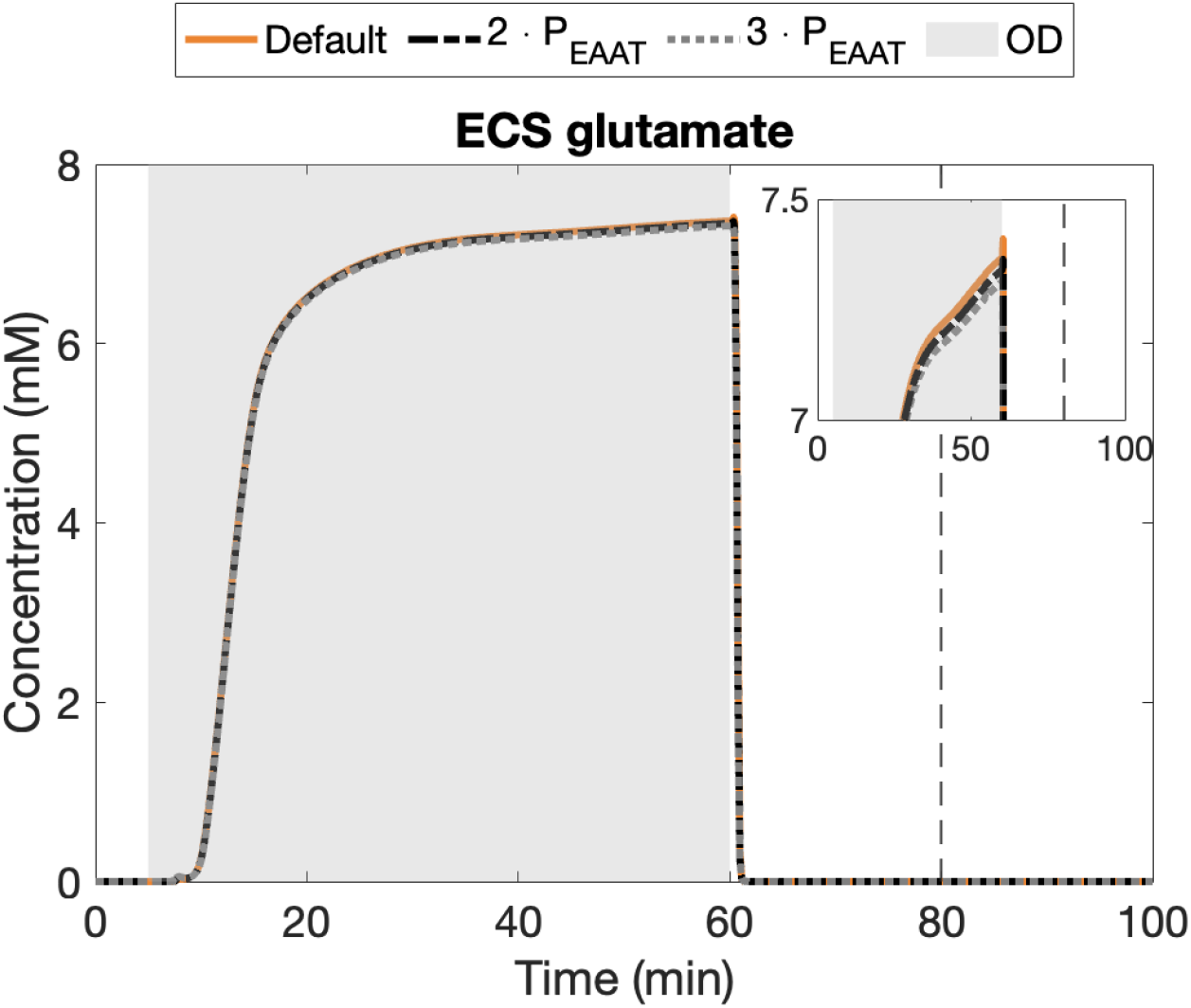
Extracellular glutamate during ischemia with increased EAAT conductance. The default simulation of ischemia (orange) is compared to simulations in which the EAAT conductance is increased twofold (black) and threefold (grey) during ischemia. Increasing the EAAT activity reduces glutamate accumulation slightly.

**Figure 8.**
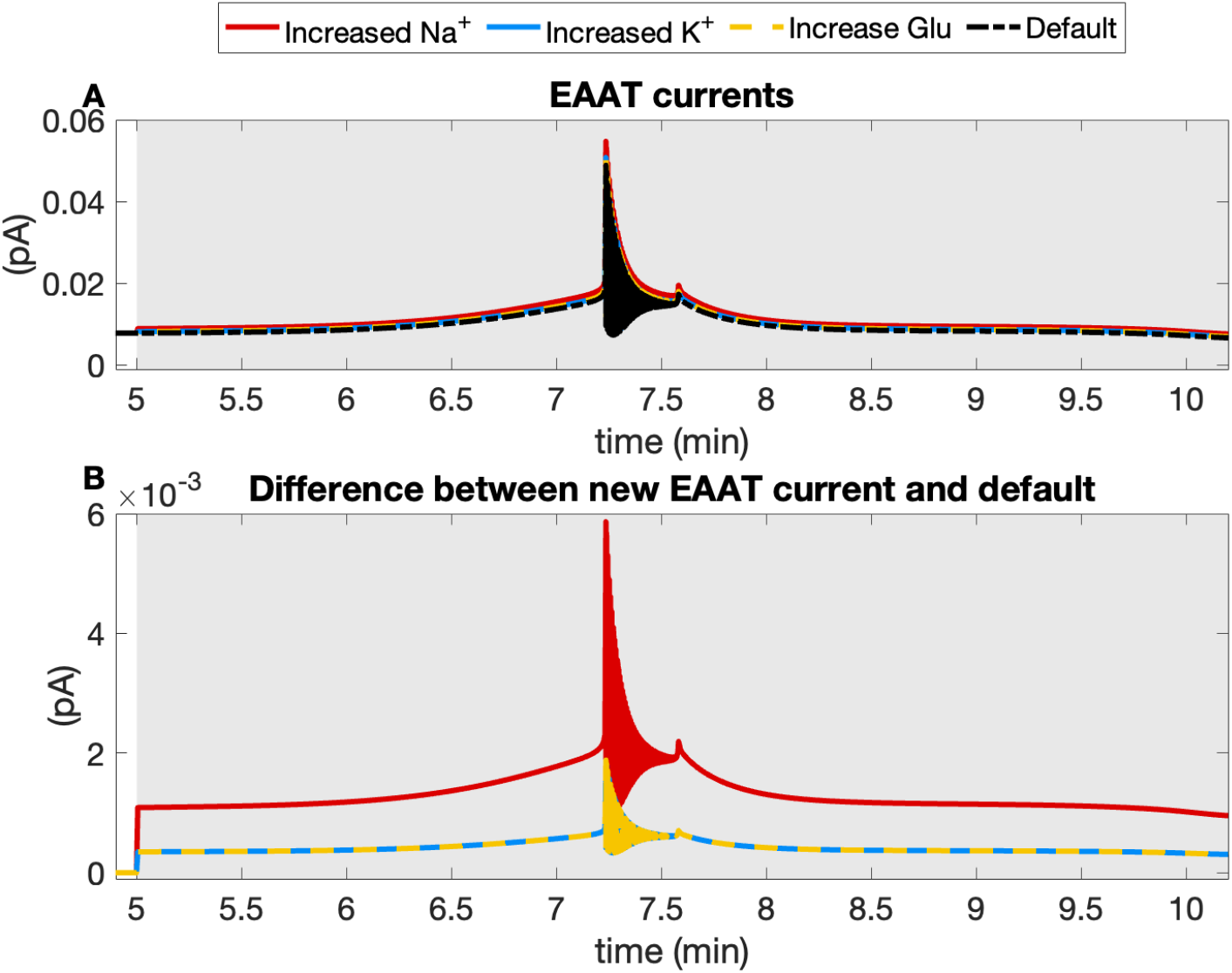
The EAAT current during ischemia. (A) The EAAT current in the default simulation of ischemia (black) is compared to EAAT currents with increased sodium (red), potassium (blue) and glutamate (yellow) gradients. The gradients are manually increased by 10% during the period of severe ischemia. (B) The difference between the altered EAAT currents due to changed gradients and the default EAAT current is shown. Modulation of potassium and glutamate gradients exerts minimal influence on the EAAT current, in contrast to the pronounced effect observed due to changes in the sodium gradient.

To determine the main factor of insufficient clearance, we analyze the three driving forces behind the EAAT: the sodium, potassium and glutamate gradients. We increase each gradient individually by 10% to assess their influence on the EAAT current. Figure 7A shows the default EAAT current alongside the modified EAAT currents resulting from altered ion gradients during ischemia, while Figure 7B presents the absolute difference between the modified and default EAAT currents. Modifying the potassium or glutamate gradient has minimal impact on the EAAT current, whereas altering the sodium gradient significantly affects the EAAT current, which is related to its 3:1:1 stoichiometry. These findings indicate that insufficient glutamate clearance, governed by the breakdown of the astrocytic sodium gradient, is the primary cause of glutamate accumulation.

### 2.3 Synaptic recovery

In addition to investigating the cause of glutamate accumulation during ischemia, we also examine which mechanisms can contribute to the recovery of synaptic transmission and restoration of the glutamate–glutamine cycle. We study three possible recovery mechanisms: increasing the glutamine synthetase activity, increasing the EAAT activity, and blocking the voltage-gated ion channels.

#### 2.3.1 Increased glutamine synthetase activity

In Figure 9, we show the effect of increasing the glutamine synthetase reaction speed. The astrocytic glutamate concentration decreases while all glutamine concentrations increase, but none return to their pre-ischemic levels. Extracellular glutamate is able to decrease to 70 nM but does not fully return to its baseline concentration of 13 nM. At t=100 minutes, we apply external stimulation to the neuron to test glutamate release. Although extracellular glutamate is partially cleared, synaptic glutamate release remains impaired.

**Figure 9.**
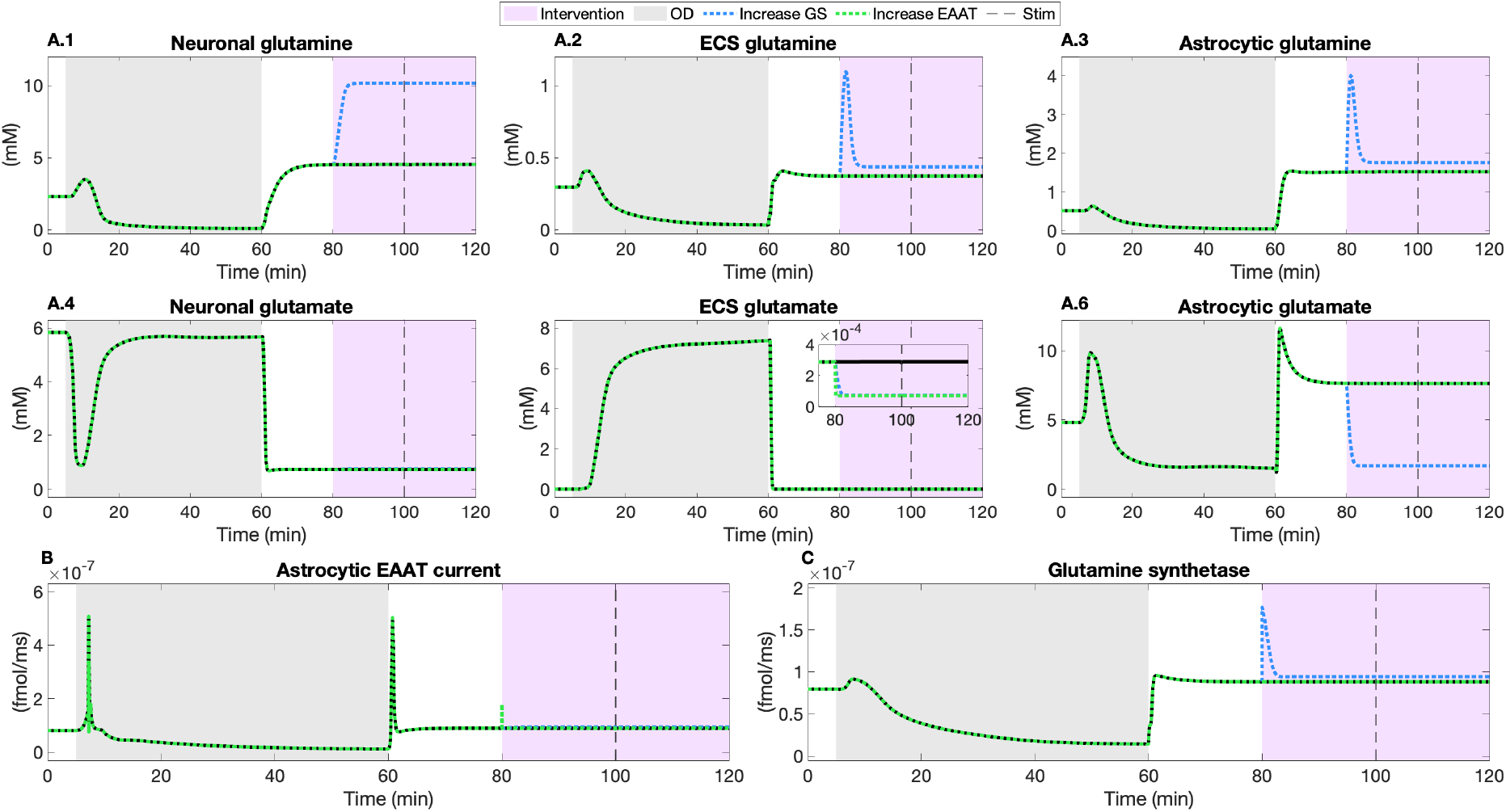
Possible recovery mechanisms after ischemia. Severe ischemia occurs from t=5 to t=60 minutes. At t=80, the activity of glutamine synthetase (blue) and the neuronal and astrocytic EAAT currents (green) are increased. Increasing the glutamine synthetase activity leads to higher glutamine concentrations, and reduced astrocytic and extracellular glutamate. Increasing the EAAT currents results in an marginal increase in glutamine concentrations, and reduced extracellular glutamate. At t=100 minutes, the neuron is stimulated but no glutamate is released in both cases. Neither of the interventions results in recovery of synaptic transmission.

#### 2.3.2 Increased astrocytic EAAT activity

Increasing the EAAT current has less impact on the glutamate and glutamine concentrations. The astrocytic glutamate concentration and all glutamate concentrations increase marginally but do not return to pre-ischemic levels, see Figure 9. Extracellular glutamate decreases, but again no glutamate is released during neuronal stimulation. Our findings indicate that even improved glutamate clearance is insufficient for recovery of synaptic transmission after ischemia.

#### 2.3.3 Blockade of voltage-gated ion channels

The reduced sodium gradient is the main factor involved in the malfunction of the EAAT and synaptic transmission failure. We hypothesized that blockade of the TTX-sensitive voltage-gated sodium channels might rescue the neuron and astrocyte from their pathological equilibrium. The neuronal sodium channel is blocked to prevent sodium influx from t=80 to t=90 minutes, shown in Figure 10. In this scenario, the neuronal NKA restores the sodium and other ion gradients, membrane potentials return to baseline values, and extracellular oxygen concentrations are restored. Due to the sodium dependence of glutamate uptake and glutamine transport, all glutamate and glutamine concentrations return to their pre-ischemic values. In response to an electrical current, the generation of an action potential activates the calcium-dependent glutamate cycle and glutamate is released into the extracellular space. All together, our results show that impaired synaptic transmission during ischemia thus results from EAAT dysfunction, which is governed by the sodium gradient. If normoxia is restored, recovery of the sodium gradient by inhibition of the voltage-gated sodium channel is sufficient for recovery of synaptic transmission.

**Figure 10.**
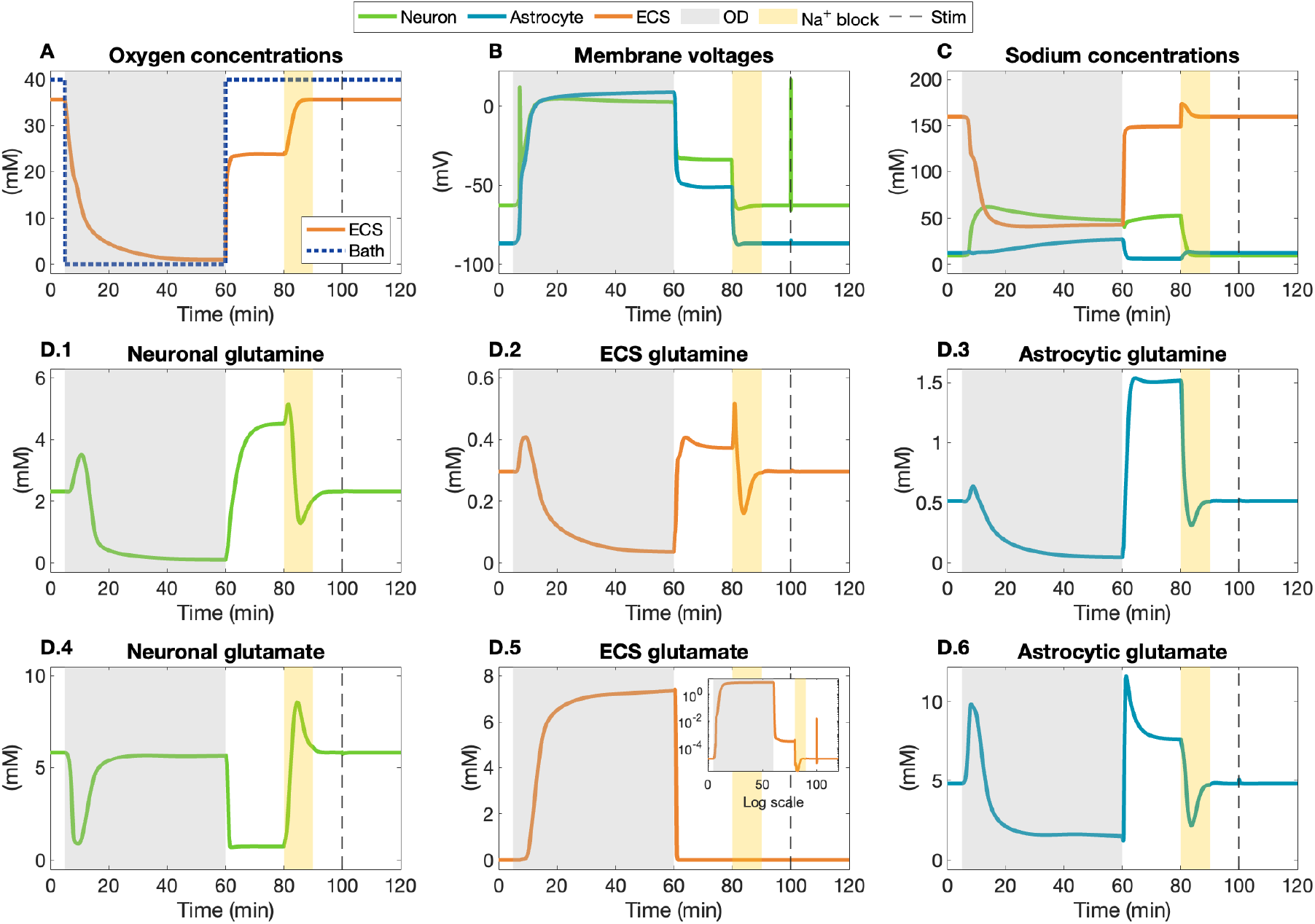
Sodium block after ischemia. Severe ischemia occurs between t=5 and t=60 minutes. From t=80 to t=90 min, voltage gated sodium channels are blocked. (A) The extracellular oxygen concentration returns to baseline after the blockade. (B) Membrane potentials return to the physiological equilibrium. At t=100 min, action potential are generated in response to external stimulation. (C) Neuronal sodium can recover due the NKA and blocked sodium influx. (D) All glutamate and glutamine concentrations return to baseline during the sodium channel block. At t=70 min, the neuron is stimulated and glutamate is released into the extracellular space. Blockade of the neuronal sodium channels results in recovery of ion homeostasis and synaptic transmission.

## 3 Discussion

In this study, we present the first detailed model of a neuron and astrocyte incorporating the complete glutamate-glutamine cycle to investigate the GG-cycle during ischemia and recovery.

### 3.1 The glutamate-glutamine cycle is essential for synaptic transmission

Our biophysical model combines ion concentrations together with the complete glutamate-glutamine cycle, which enables a detailed analysis of synaptic transmission during ischemia. Implementation of the glutamate-glutamine cycle is essential to study synaptic transmission as it provides the main pathway for astrocytic glutamate back to the neuron (Flores-Méndez et al., 2016; Nicoli et al., 2024). Experimental studies demonstrate that inhibiting glutamine synthetase reduces glutamate release and thus impairs synaptic transmission (Rothstein & Tabakoff, 1984; Tani et al., 2014), as we verify in Figure 1. Therefore, incorporating the essential GG-cycle when studying synaptic transmission during ischemia seems warranted.

### 3.2 Severe ischemia leads to disturbed ion homeostasis and glutamate accumulation

While moderate ischemia induces only transient changes in ion and glutamate concentrations, severe ischemia has significant and irreversible pathological consequences. The differences in neuronal and synaptic damage between moderate and severe ischemia are also observed experimentally (Ueda et al., 1992; Winkelheide et al., 2008; Zhao et al., 1998). Severe ischemia leads to a significant disturbance in ion homeostasis, as shown in Figure 3. Strongly increased intracellular sodium due to decreased NKA activity is one of the primary consequences of severe ischemia (Gerkau et al., 2017; van Putten et al., 2021). The extracellular potassium concentration increases due to decreased NKA activity, also determined experimentally (Kléber, 1984). Furthermore, the simulations show highly elevated intracellular calcium concentrations, similar to elevated calcium levels observed experimentally (Jalini et al., 2016; Silver & Erecińska, 1990), which affects calcium-dependent glutamate release. In conclusion, the dynamics of ion concentrations affecting synaptic transmission align with experimental findings.

Besides disturbed ion homeostasis, an important consequence of severe ischemia is glutamate accumulation in the extracellular space. Glutamate accumulation in the extracellular space occurs in two distinct phases, as observed experimentally (Ueda et al., 1992; Wahl et al., 1994). At the onset of ischemia, the initial phase is marked by a moderate rise in extracellular glutamate levels due to neuronal firing. The second phase occurs during the period of neuronal depolarization and depression, in which extracellular glutamate increases substantially, up to the millimolar range. The two-stage glutamate accumulation observed in our study is also reported by (Ziebarth et al., 2025). Besides glutamate accumulation, we show that no glutamate is released upon additional neuronal stimulation, indicating synaptic transmission failure.

### 3.3 The main cause of glutamate accumulation is disrupted uptake

While the exact cause of synaptic transmission failure is not yet fully understood, experimental data suggest three plausible causes: decreased activity of energy-dependent glutamine synthetase, excessive glutamate release by neurons and astrocytes and reduced glutamate uptake or reverse uptake by the EAATs.

First, an experimental simulation in which glutamine synthetase is independent of energy shows that increased glutamine synthetase activity results in increased glutamine concentrations. The surplus of glutamine is converted into glutamate, which accumulates in the ECS. Altering glutamine synthetase to be independent of energy thus aggravates glutamate accumulation, suggesting that its dysfunction is unlikely to be the main cause of glutamate accumulation.

Subsequently, we demonstrate that the extracellular glutamate concentration only increases marginally due to glutamate release during repetitive neuronal firing in the first several minutes of ischemia. The majority of glutamate accumulation occurs during the remaining phase of ischemia, in which neuronal activity is suppressed. Furthermore, our results show that glutamate accumulation is not aggravated by reversed glutamate uptake, as is proposed in (Grewer et al., 2008; Rossi et al., 2000). The reversal potential of the EAAT is substantially more positive than the membrane potentials, see Figure 16. Thus, excessive glutamate release is excluded as the primary cause of glutamate accumulation.

As we have eliminated glutamate release and malfunction of the GG-cycle as possible causes, the remaining potential cause is a reduction of the astrocytic EAAT current. Our findings are consistent with experimental evidence indicating that, under ischemic conditions, glutamate uptake by the EAAT is significantly limited (Bruhn et al., 2001). Our results indicate that EAAT function is predominantly governed by the sodium gradient, with minimal contribution from the potassium and glutamate gradients, as shown experimentally (Kelly et al., 2009; Rose et al., 2018). Our research not only verifies reduced glutamate uptake by the EAAT transporter during ischemia, but also identifies it as the main contributor to glutamate accumulation in the ECS.

### 3.4 Recovery of synaptic transmission is possible by restoring the sodium gradient

We have shown that an increase in either glutamine synthetase activity or EAAT activity leads to reduction of the extracellular glutamate concentration. However, none of these interventions results in recovery of synaptic glutamate release. We therefore conclude that the modifications to the GG-cycle are insufficient as a recovery mechanism. Since EAATs malfunction due to an altered sodium gradient, we investigate restoring the sodium gradient as a possible recovery mechanism. Indeed, blocking voltage-gated TTX-sensitive neuronal sodium channels results in recovery of ion homeostasis, and in turn, recovery of synaptic transmission.

### 3.5 Limitations

There are four main limitations of our model. Firstly, there are no spatial components, meaning that we consider each compartment to be uniform and do not include spatial diffusion. As we consider only one synapse in an ECS, there is no ion or neurotransmitter spillover to other synapses. Due to the fixed total volume of the model, there is also no ion diffusion due to (re)perfusion such as in an experimental set-up. As a result, ionic changes during prolonged ischemia in the brain could be even more pronounced than those observed in experiments. However, given that this modeling assumption impacts all ionic gradients uniformly, it does not compromise our main conclusions.

Secondly, the SN transporter, responsible for glutamine transport at the astrocytic membrane, is not able to reverse in our model. While experimental studies show that SN reversal can occur in response to pH changes (Todd et al., 2017), our model does not include proton dynamics and therefore does not account for this reversal mechanism. The most important driving force for the SN transporter is the sodium influx by the EAAT (Todd et al., 2017), and changes in the proton gradient are unlikely to have a dominant impact. Furthermore, intracellular and extracellular compartments both acidify similarly during ischemia, largely preserving the pH gradient (Hagberg, 1985; von Hanwehr et al., 1986).

Thirdly, we consider only one calcium-compartment in the neuron and astrocyte. A differentiation between cytosolic and the endoplasmic reticulum (ER) would introduce more detailed dynamics and timescales. It has been shown that calcium dynamics operate on several timescales in the cytosolic and ER (Bazargani & Attwell, 2016; Cheng et al., 2006). During ischemia, the cytosolic calcium concentration increases partly due to calcium release from intracellular stores like the ER (Bodalia et al., 2013; Paschen & Doutheil, 1999). The calcium compartment in our model represents the cytosol, and experiences a large increase in concentration during ischemia. While the cause of increased calcium might differ in vivo, we argue that the resulting effect of elevated calcium is similar. For future work, it would be interesting in investigating if the difference between cytosol calcium and ER stores influences the timescale and extent of synaptic transmission failure.

Lastly, our model does not include a postsynaptic neuron. In order to fully investigate synaptic transmission and glutamate toxicity in ischemia, it is necessary to analyze postsynaptic glutamate receptors such as NMDA and AMPA receptors. These receptors are overstimulated during ischemia, thereby releasing potassium into the ECS, and they might desensitize during glutamate toxicity (Olloquequi et al., 2018; Sattler & Tymianski, 2001). Our simulations show an extracellular glutamate concentration of around 7.5 mM during ischemia. Due to the lack of spatial diffusion and the postsynaptic neuron, this concentration is likely to be higher than seen in experiments. However, glutamate toxicity is already observed for much lower extracellular glutamate concentrations (Michaels & Rothman, 1990), thus we expect that postsynaptic receptors would be overstimulated in our simulation.

In conclusion, we have constructed a detailed model that includes the first implementation of the complete glutamate-glutamine cycle. With the model, we simulate different severities of ischemia, to investigate the primary cause of synaptic transmission failure and glutamate accumulation. We demonstrate that during severe ischemia, characterized by a transient complete cessation of energy supply, extracellular glutamate levels can rise to the millimolar range, sufficient to induce glutamate toxicity. Our findings show that the main cause of glutamate accumulation during severe ischemia is the reduced activity of the astrocytic EAAT due to the degraded sodium gradients. Furthermore, we demonstrate that alterations to the GG-cycle are not sufficient to mediate recovery, whereas blocking the neuronal sodium channels restores synaptic transmission.

## 4 Methods

We elaborate on the model of (Kalia et al., 2021) and consider a glutamatergic presynaptic neuron and an associated astrocyte confined in a finite extracellular space, shown in Fig 11. The ECS is surrounded by an oxygen bath that allows oxygen diffusion to the ECS. In this model, oxygen serves as a proxy for ATP and is considered to be the energy source. Transmembrane ion fluxes for sodium, potassium, and chloride are described by voltage-gated channels and leak channels. To maintain ion homeostasis, the transmembrane ion fluxes are counteracted by ion transporters, such as the energy-dependent sodium-potassium ATPase, the potassium-chloride transporter (KCC), and the sodium-potassium-chloride cotransporter (NKCC1). Volume regulation is based on differences in osmotic pressure, which can lead to cell swelling. Calcium dynamics in the synaptic compartments are described by voltage-gated and leak channels, supported by the sodium-calcium exchanger (NCX). To simulate stimulation experiments, a square-wave sodium current is applied, used for action potential generation. To model synaptic transmission, we introduce the first complete overview of glutamate-glutamine dynamics. For glutamate endocytosis and exocytosis, we combine calcium-dependent glutamate release (Fig 12) with uptake by the excitatory amino acid transporter (EAAT). To complete glutamate recycling, we implement the glutamate-glutamine cycle, depicted in Fig 13. In this cycle, astrocytic glutamate is converted into glutamine by the ATP-dependent enzyme glutamine synthetase (Petito et al., 1992; Schousboe et al., 2014). Subsequently, glutamine is transported via ECS back to the neuron by the SN and SAT transporters (A. Bröer et al., 2002; S. Bröer & Brookes, 2001; Chaudhry et al., 2002; Mackenzie & Erickson, 2004; Todd et al., 2017; Yao et al., 2000). Finally, glutamine is converted back into glutamate by the enzyme glutaminase and glutamate moves through the calcium-dependent glutamate cycle (Bak et al., 2006; Kalia et al., 2021; Krebs, 1935), where it is stored in the neuronal depot. Ischemia is simulated by lowering the oxygen concentration in the bath manually. Due to unrestricted exchange between the bath and the ECS, this results in restricted energy supply for the neuron and astrocyte. Different severities of ischemia are simulated by changing the duration of reduced oxygen and thereby the level of oxygen in the bath.

**Figure 11.**
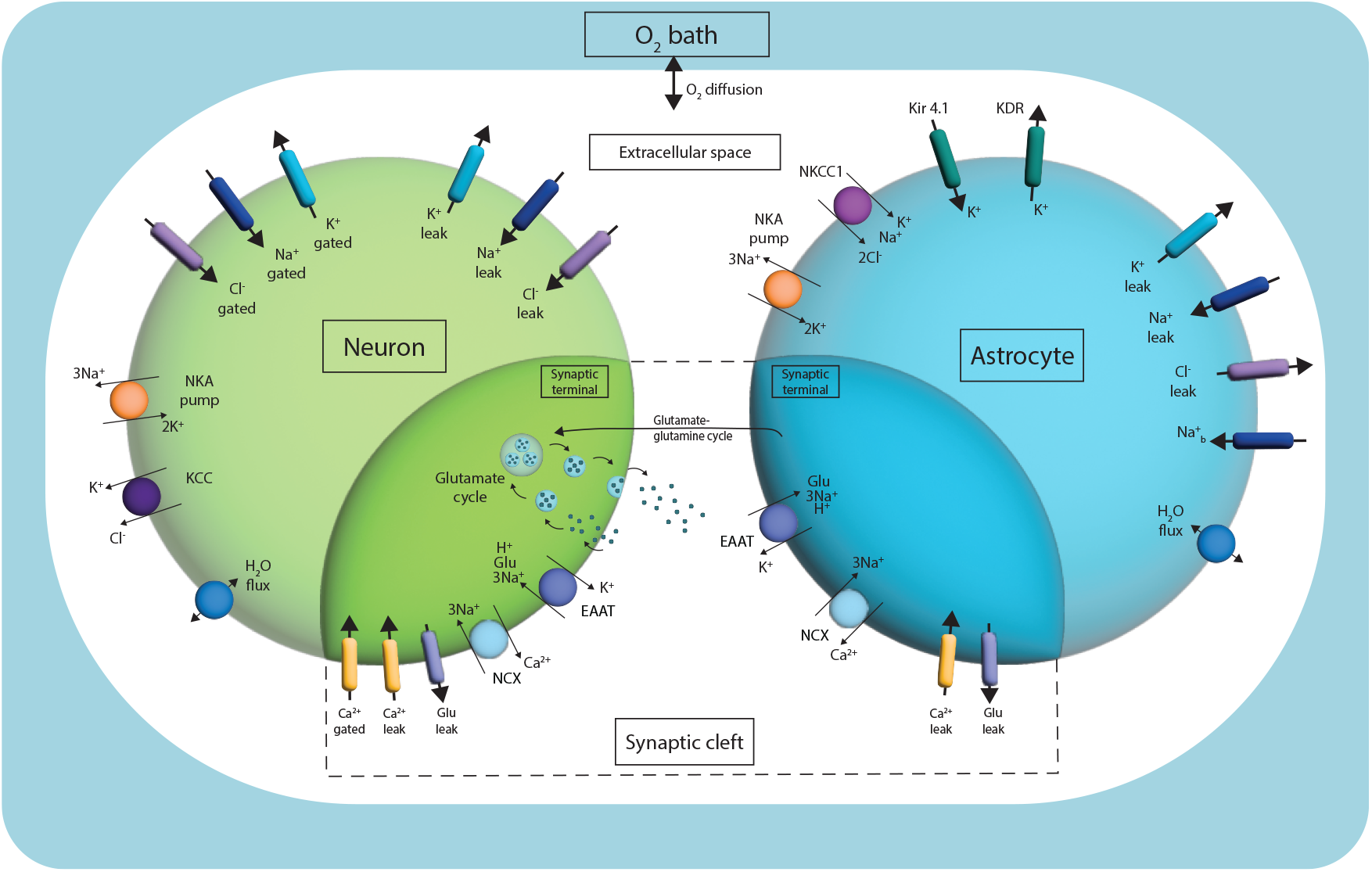
Schematic overview of the model. The model includes six compartments: three main compartments for the neuron, astrocyte and extracellular space and three synaptic compartments: the neuronal and astrocytic synaptic terminal and the synaptic cleft. In the main compartments, sodium, potassium and chloride dynamics are considered. In the synaptic compartments, calcium, glutamate and glutamine concentrations are additionally included. Glutamate and glutamine dynamics in the astrocyte and neuron are described by the glutamate-glutamine cycle. The extracellular space is surrounded by an oxygen bath that allows oxygen diffusion between the bath and extracellular space. The main ion transporters are the energy-dependent Na^+^/K^+^-ATPase (NKA) and the Na^+^-dependent glutamate transporter excitatory amino acid transporter (EAAT), located in the main compartment and synaptic compartment, respectively. KCC: K^+^/Cl^−^-cotransporter, NKCC1: Na^+^/K^+^/Cl^−^-cotransporter, Kir4.1: K^+^ inward rectifier channel 4.1, NCX: Na^+^/Ca^2+^-cotransporter. KDR: K^+^ delayed rectifying current.

**Figure 12.**
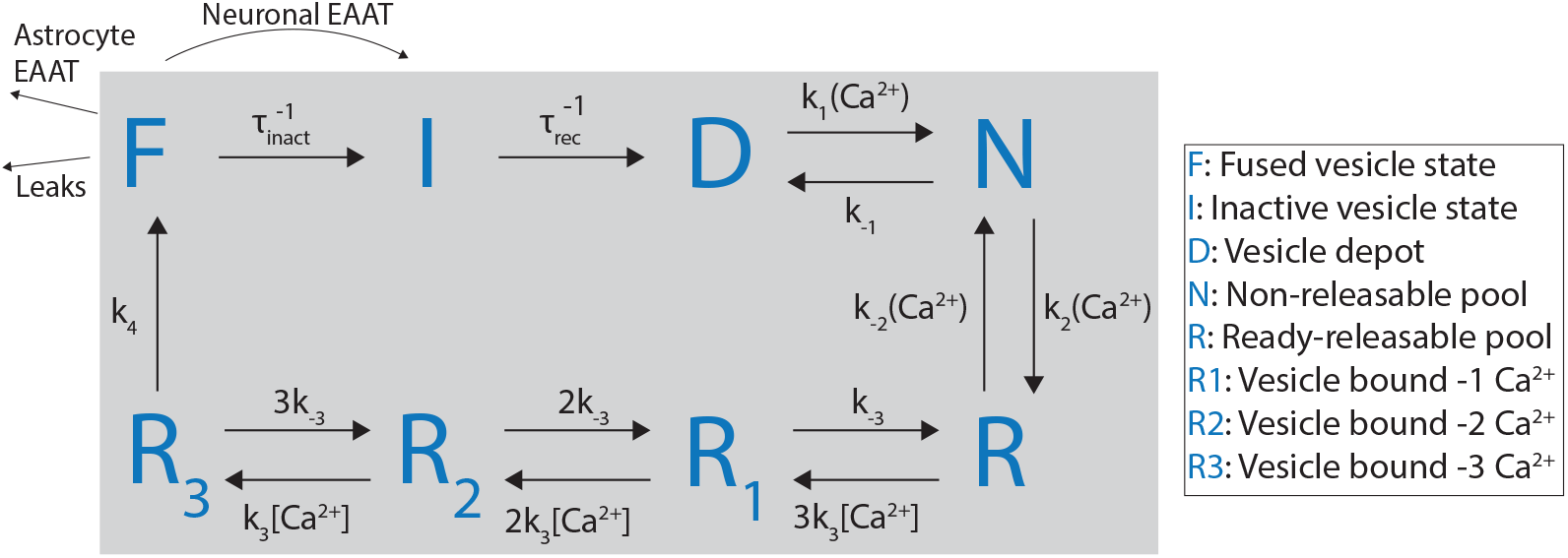
Overview of the glutamate cycle from (Kalia et al., 2021). The cycle consists of 8 states with reaction rates *τ*_*rec*_, *k*_*i*_, which are forward or backward rates and can be calcium-dependent. Starting at the glutamate depot, where most neuronal glutamate is located, glutamate moves through five calcium-dependent states (N, R, R1, R2, R3) before it is released into the extracellular space. From the fused state, the EAAT transports glutamate into the first state in the neuron, the inactive vesicle state. From the inactive state, glutamate is packed into vesicles and transported to the depot.

**Figure 13.**
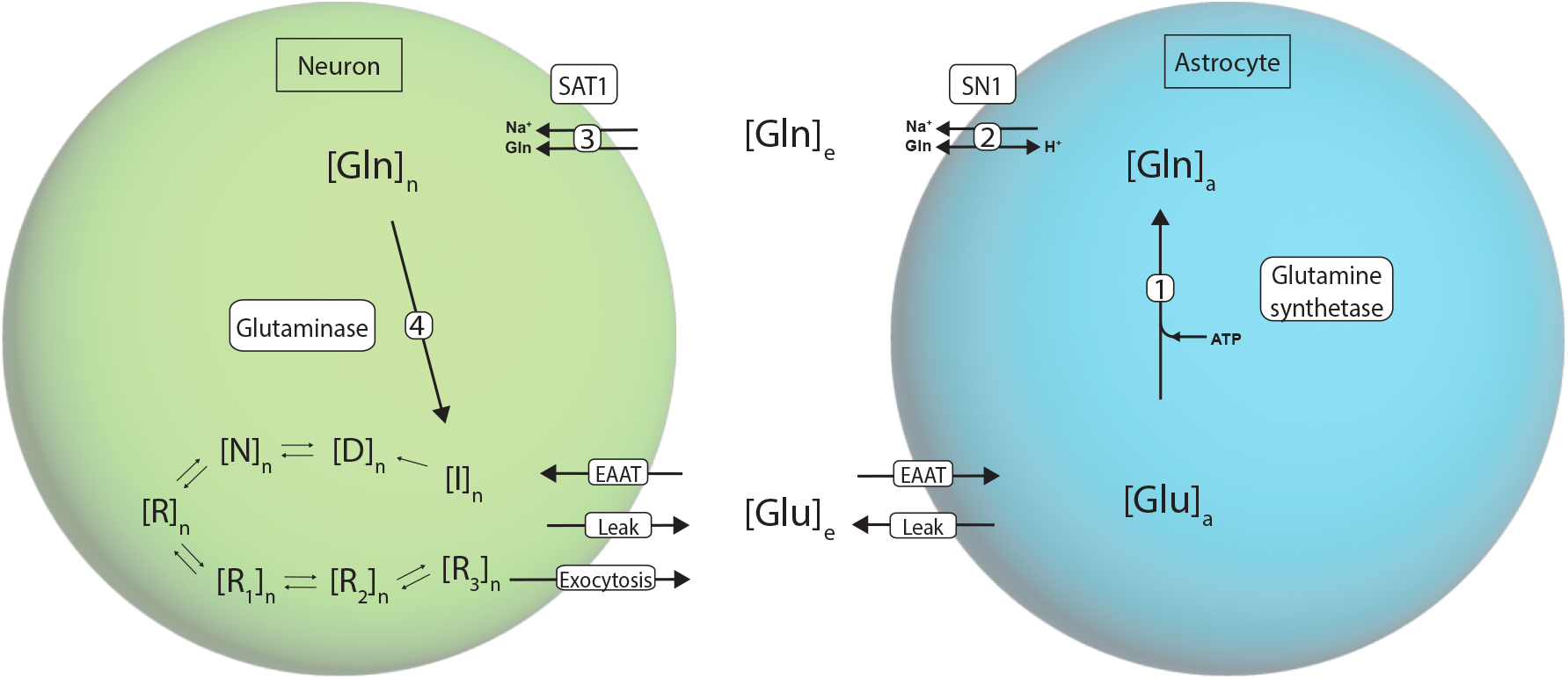
Overview of the glutamate-glutamine cycle and the glutamate cycle. The glutamate-glutamine cycle consists of four steps. First, astrocytic glutamate is converted into glutamine by the enzyme glutamine synthetase, an ATP-dependent step. Second, astrocytic glutamine along with sodium is transported to the extracellular space by the system N transporter (SN1). Third, from the extracellular space, sodium and glutamine are transported into the neuron by the system A transporter (SAT1). Lastly, neuronal glutamine is converted back into glutamate by the enzyme glutaminase, and transported to the glutamate cycle, specifically the inactive state. From here, glutamate can move through all states of the glutamate cycle before it is excreted to the extracellular space.

### 4.1 The model equations

The model equations are based on the rate of change of the molar amount for each ion, which is determined by the sum of all currents. The gating variables for the voltage-gated sodium, potassium and chloride channels are based on the Hodgkin-Huxley formalism. Lastly, volume dynamics depend on the difference in osmolarity, i.e., the difference in total concentrations in each compartment. The fundamental model equations are:

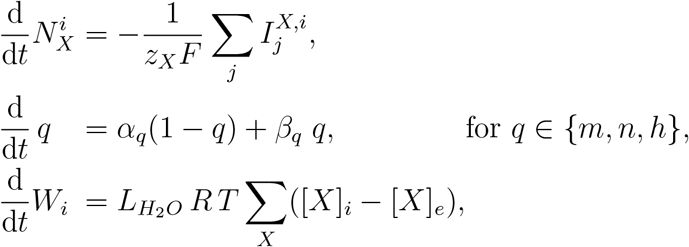

where *z*_*X*_ is the valency of ion *X, F* is Faraday’s constant, 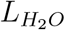 is the neuronal membrane water permeability and *R* is the universal gas constant. The number of moles for ion *X* in compartment *i* is denoted by 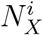. The volume of compartment *i* is *W*_*i*_. Subsequently, the concentration of ion *X* in compartment *i* is computed as 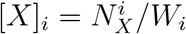. The rate of change of the number of moles is the sum of all currents 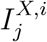 affecting ion species *X*, where *j* denotes the type of current. These fundamental equations are extended with the glutamate cycle and the glutamate-glutamine cycle explained in further sections.

The model adheres to three conservation laws: the conservation of charge, volume and mass, which are formulated as follows

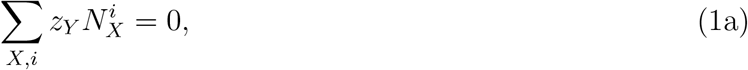

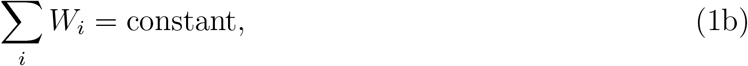

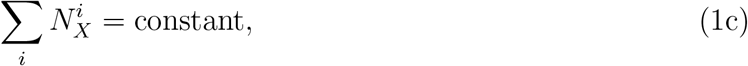

where the conservation of charge and mass are formulated for each ion *X*. The conservation laws are used to compute the extracellular volume and molar amounts as follows,

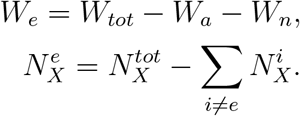

### 4.2 Oxygen dynamics

We have adapted the oxygen dynamics from (Wei et al., 2014). The extracellular space is enclosed by a bath with a constant oxygen concentration, enabling oxygen diffusion to the extracellular space. The model is expanded with the following model equation,

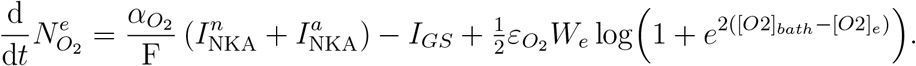

### 4.3 The glutamate cycle

The glutamate cycle concerns the presynaptic vesicle formation, docking and exocytosis. Upon arrival of an action potential in the presynaptic neuron, the voltage-gated calcium channels open, leading to a large influx of calcium. The influx of calcium phosphorylates the protein synapsin I, responsible for immobilizing vesicles (Nichols et al., 1992). Phosphorylated synapsin releases vesicles from the depot, enabling vesicles to move towards the membrane, where the SNARE protein is located. Upon calcium binding to synaptotagmin, the SNARE protein is activated and vesicles can fuse with the membrane, resulting in exocytosis of glutamate (Nichols et al., 1992). After exocytosis, approximately 10% of glutamate is taken up by EAATs into the neuron, where it is packed into vesicles by vesicular glutamate transporters (VGLUTs) and will be stored in the vesicle depot. This cycle is captured in eight states as proposed by (Kalia et al., 2021) who combined the work in (Tsodyks & Markram, 1997; Walter et al., 2013). The overview of all states with corresponding kinetic rates is shown in Figure 12. The model equations for the glutamate cycle can be found in in the Appendix.

### 4.4 The glutamate-glutamine cycle

The other 90% of glutamate is taken up by the astrocytic EAAT and passes through the glutamate-glutamine cycle, which is essential for glutamate homeostasis (Bak et al., 2006; Berl & Clarke, 1983). We consider four steps in the glutamate-glutamine cycle, which are shown in Figure 13. First, astrocytic glutamate is converted into glutamine by the enzyme glutamine synthetase. Glutamate synthesis is ATP-dependent and will not function properly during metabolic stress (Passlick et al., 2021). After the synthesis of glutamate, glutamine is transported to the neuron via a two-step process by system A and system N transporters. We consider the transporters SN1 and SAT1, also known as SNAT3 and SNAT1, respectively, and reviewed in (Mackenzie & Erickson, 2004). The electroneutral SN1 transporter controls glutamine efflux from the astrocyte and transports sodium alongside glutamine out of the astrocyte in exchange for a proton (Chaudhry et al., 2002). This transporter is known to be pH sensitive and has been demonstrated to reverse direction at low pH levels (Chaudhry et al., 1999; Chaudhry et al., 2002). Subsequently, the SAT1 transporter takes up glutamate alongside sodium from the ECS into the neuron (Chaudhry et al., 2002). Finally, neuronal glutamine is converted to glutamate by the enzyme glutaminase (Bak et al., 2006), first reported by Krebs (Krebs, 1935). In conclusion, the glutamate-glutamine cycle consists of four steps, including energy-dependent glutamine synthesis.

#### 4.4.1 The glutamate-glutamine cycle model equations

The dynamics of all four steps are described using Michaelis-Menten (MM) kinetics. The MM-constants for each step are extracted from experimental studies (Bak et al., 2006; Kvamme et al., 2001; Mackenzie et al., 2003; Schousboe et al., 2014; Todd et al., 2017; Uwechue et al., 2012; Yao et al., 2000). We note that the in the first step, the enzyme glutamine synthetase requires ammonium. Ammonium is a byproduct of the glutamate synthesis in the neuron (step 4). It is suggested that an ammonium pathway exists, along which ammonium produced in the neuron is transported back to the astrocyte where it is used for glutamine synthesis (Bak et al., 2006; Lieth et al., 2001; Waagepetersen et al., 2000). Therefore, we assume a constant ammonium concentration that is sufficient for optimal function of glutamine synthetase. Similarly, we assume a constant pH and thus a constant proton concentration in the astrocyte and the extracellular space. The full model equations of the GG-cycle are given by,

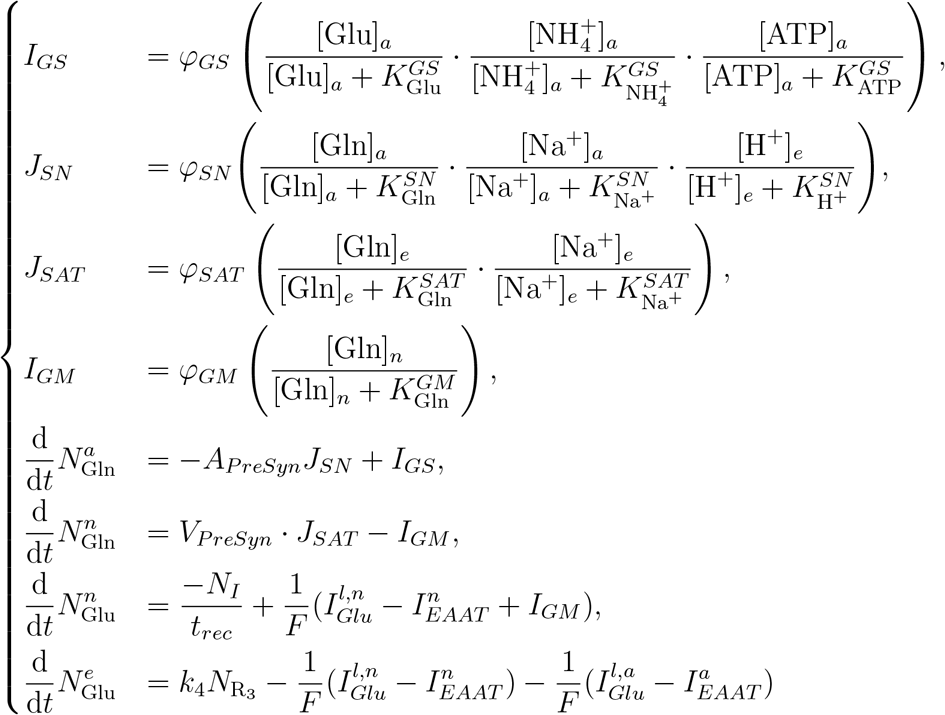

where we assume 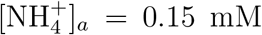, and an astrocytic pH of approximately 7.2. Further-more, 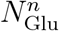 refers to the inactive state, which is the first neuronal glutamate state, see Fig 12. Extracellular glutamine follows from the conservation laws.

## A Supplementary model components

The astrocytic potassium delayed rectifying channel (KDR) was added to the model to provide an additional pathway for efflux of potassium in the astrocyte. The formulation is taken from (Dronne et al., 2006) and is given by:

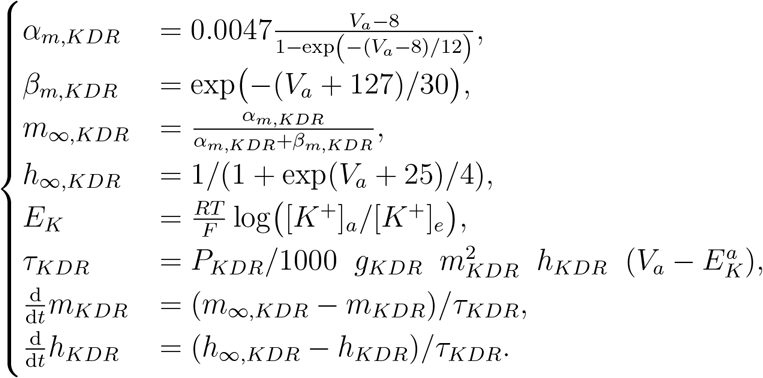

The formulation for the EAAT transporter is taken from (Breslin et al., 2018). The EAAT transporter is based on the voltage difference between the membrane potential and the reversal potential of the transporter as follows:

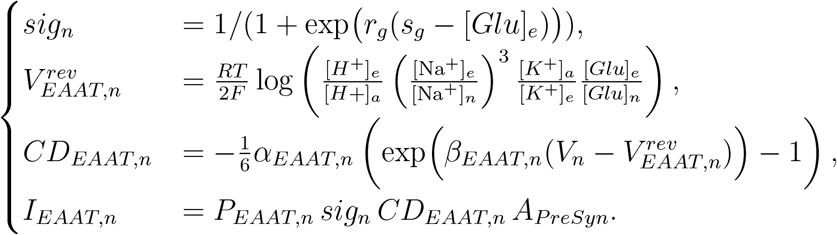

As the glutamate-glutamine cycle provides an additional astrocytic sodium efflux by the SN-transporter, an additional influx was needed. We implemented a straightforward sodium current 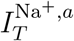 based on the Nernst potential, similar to that in (Breslin et al., 2018), with the following formulation:

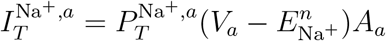

## B Parameter fitting

The parameters needed for implementation of the GG-cycle cannot be sourced from literature due to a lack of data on timescales of glutamate and glutamine dynamics. Parameter selection is performed through parameter fitting, in which a loss function comprising two components is optimized. Firstly, we construct a loss function to ensure adequate timescales, i.e., the concentrations return to baseline after stimulation in adequate time. In order to so, we choose two time points in the simulation, where we consider ten data points at both time points. The first time point, *t*_1_, is located right before the action potential. The second time point, *t*_2_, is located after the action potential. The loss function is based on the differences between the values of the state variables at both time points. To ensure that all variables are normalized, we multiply with the weight vector *W*_1_, leading to the equilibrium loss defined as:

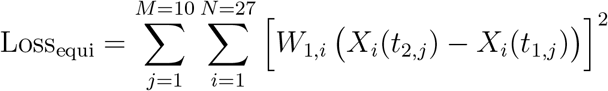

Secondly, we construct a loss function to ensure concentrations in a physiological range. To do so, we have constructed a range, denoted by *r* for each state variable, based on experimental data. In the loss function, we measure the distance of the variable to its physiological range. Subsequently, we multiply with a weight vector *W*_2_ to normalize the variables, which can also be used to prioritize certain state variables. The loss function to ensure a physiological range is defined as:

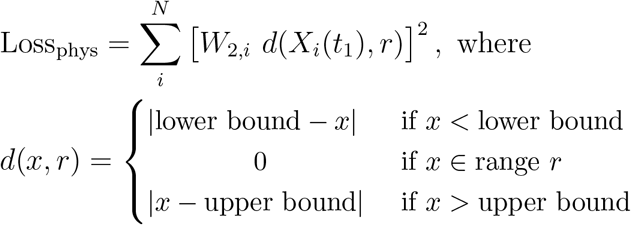

All weights can be found in Table 1.

| i | 1 | 2 | 3 | 4 | 5 | 6 | 7 | 8 | 9 | 10 | 11 | 12 | 13 | 14 |
| --- | --- | --- | --- | --- | --- | --- | --- | --- | --- | --- | --- | --- | --- | --- |
| $X_i$ | m | h | n | $N_{Na^+}^n$ | $N_{K^+}^n$ | $N_{Cl^-}^n$ | $W_n$ | $N_{Ca^{2+}}^n$ | $N_I$ | $N_D$ | $N_N$ | $N_R$ | $N_{R_1}$ | $N_{R_2}$ |
| $W_{1,i}$ | 0 | 0 | 0 | 0 | 0 | 0 | 0 | 0 | 0 | 1000 | 0 | 0 | 0 | 0 |
| $W_{2,i}$ | 0 | 0 | 0 | 0.001 | 0.001 | 0.001 | 0 | $10^7$ | 0 | 1000 | 0 | 0 | 0 | 0 |

**Table 1:**
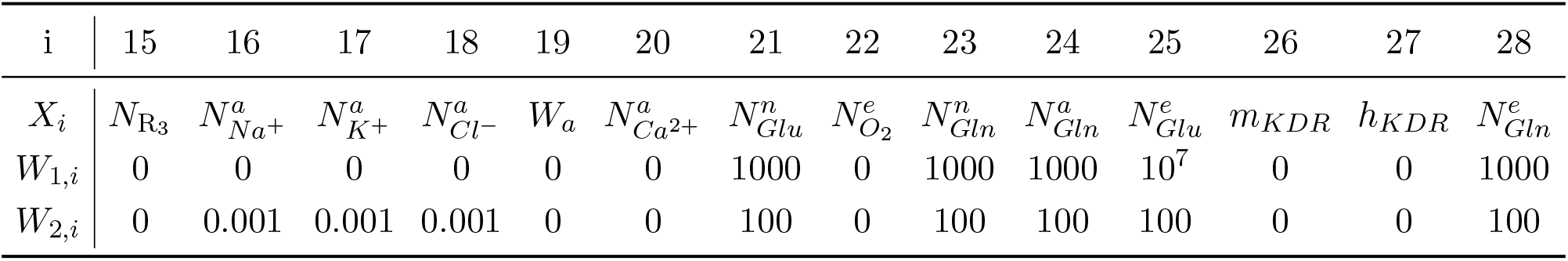
Values of weights1 and weights2.

**Table 2:**
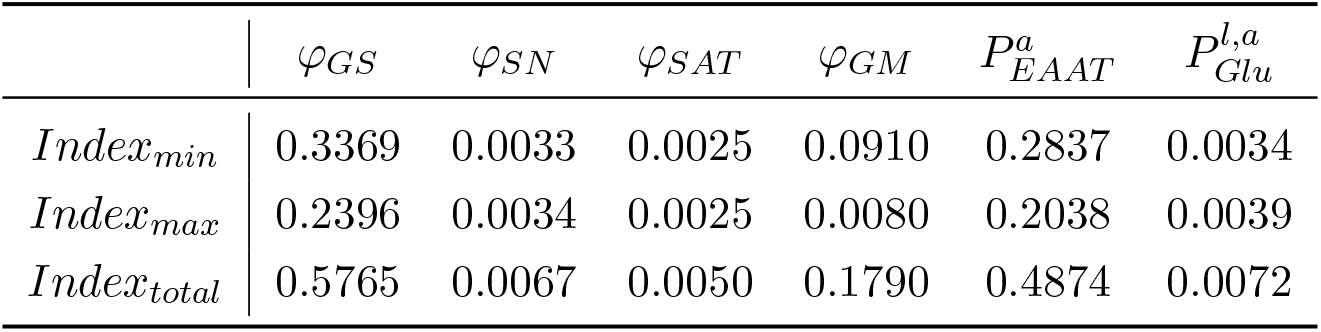
Sensitivity Indices with Min and Max Ranges.

| | $\varphi_{GS}$ | $\varphi_{SN}$ | $\varphi_{SAT}$ | $\varphi_{GM}$ | $P_{EAAT}^a$ | $P_{Glu}^{l,a}$ |
| --- | --- | --- | --- | --- | --- | --- |
| $Index_{min}$ | 0.3369 | 0.0033 | 0.0025 | 0.0910 | 0.2837 | 0.0034 |
| $Index_{max}$ | 0.2396 | 0.0034 | 0.0025 | 0.0080 | 0.2038 | 0.0039 |
| $Index_{total}$ | 0.5765 | 0.0067 | 0.0050 | 0.1790 | 0.4874 | 0.0072 |

### B.1 The optimization

In order to optimize the loss functions, the Borg Multiobjective Evolutionary Algorithm (BorgMOEA) algorithm from the global optimization Julia package BlackBoxOptim was used. The algorithm combines *ε*-dominance with Pareto solutions.

### B.2 Sensitivity analysis

After optimizing the loss functions, we performed a sensitivity analysis to evaluate the robustness and reliability of the result. For the sensitivity analysis, we use the one-factor-at-a-time method (OAT). We increase and decrease each parameter by 5%, while keeping the other parameters fixed. Then we compute the loss function again to see if changing the parameter had a large influence on the result. We compute the sensitivity index as follows:

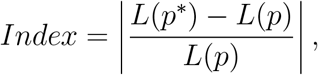

where *L*(*p*) is the loss function for parameter *p*, and *p*^*^ is the parameter vector for 5% increase or decrease.

From the sensitivity indexes, we conclude that *φ*_*GS*_ affects the result the most, but all sensitivity indexes are sufficiently low. However, changing *φ*_*GS*_ results in a slightly different equilibrium but does not affect the timescale of the model, as can be seen in Figure 14. We conclude that the results obtained from parameter fitting are reliable and robust.

**Figure 14.**
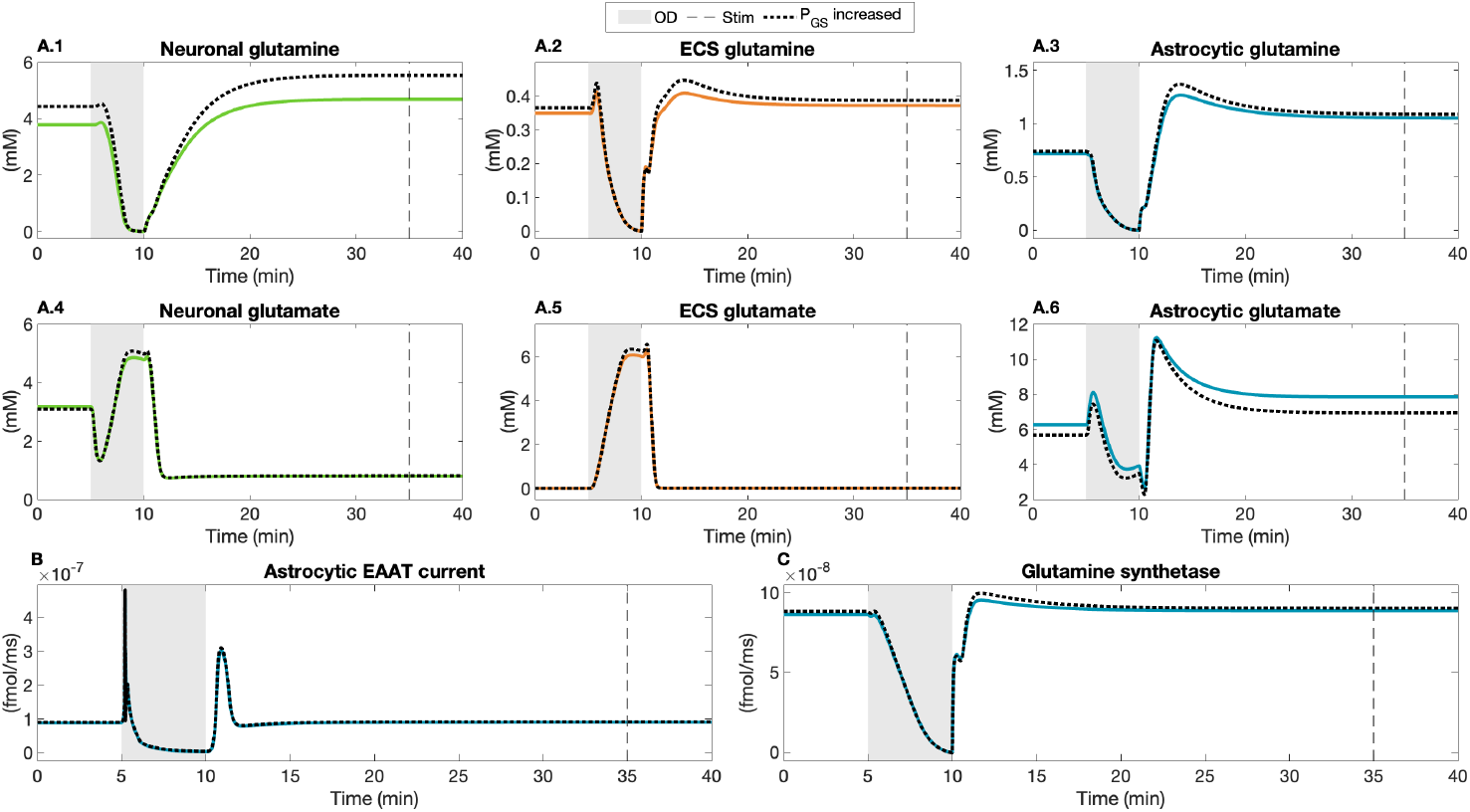
parison between default simulation of severe ischemia, and altered simulation with *φ*_*GS*_ increased by 5%.

## C Additional figures

**Figure 15.**
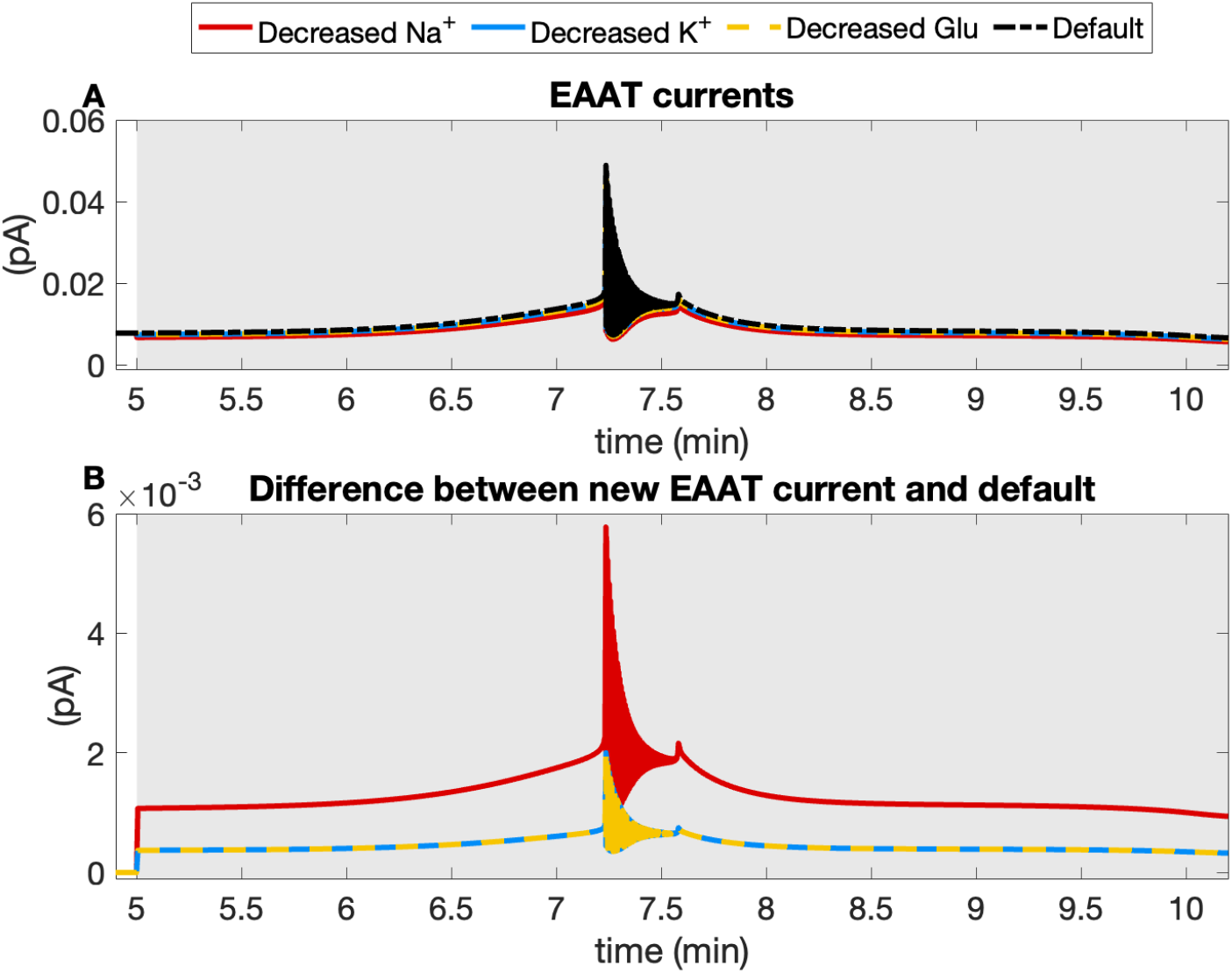
The EAAT current during ischemia. (A) The EAAT current in the default simulation of ischemia (black) is compared to EAAT currents with decreased sodium (red), potassium (blue) and glutamate (yellow) gradients. The gradients are manually decreased by 10% during the period of severe ischemia. (B) The difference between the new EAAT currents due to changed gradients and the default EAAT current is shown.

**Figure 16.**
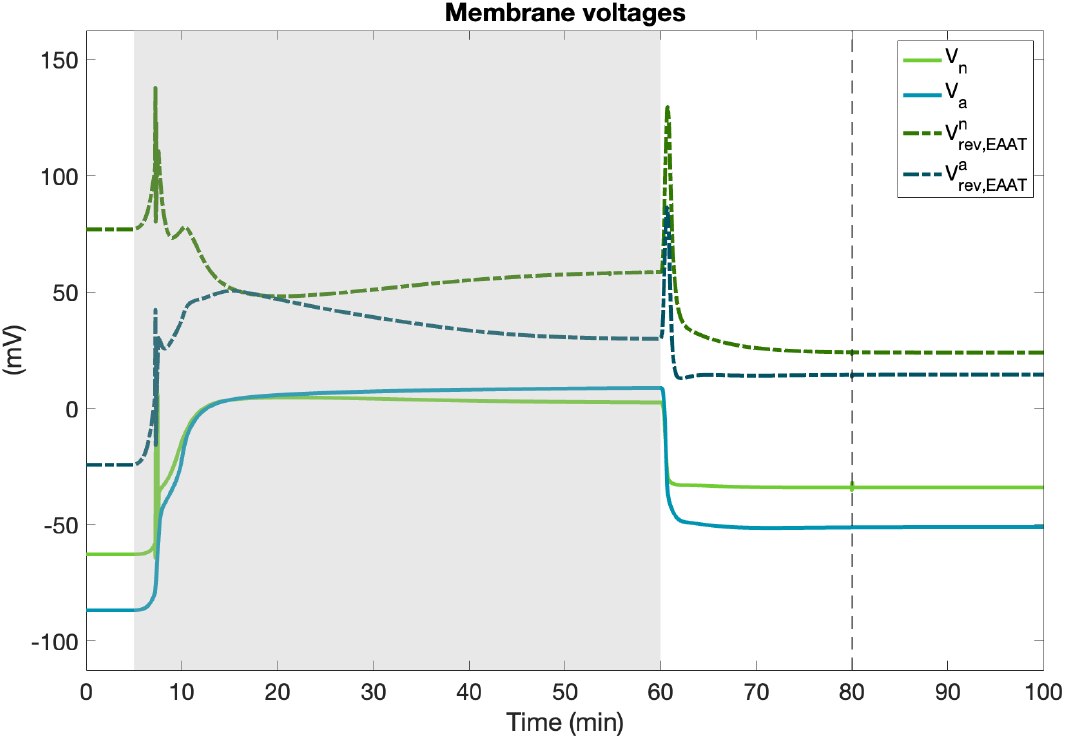
Membrane potentials and EAAT reversal potentials during ischemia.

## D The complete model equations

### D.1 Neuron

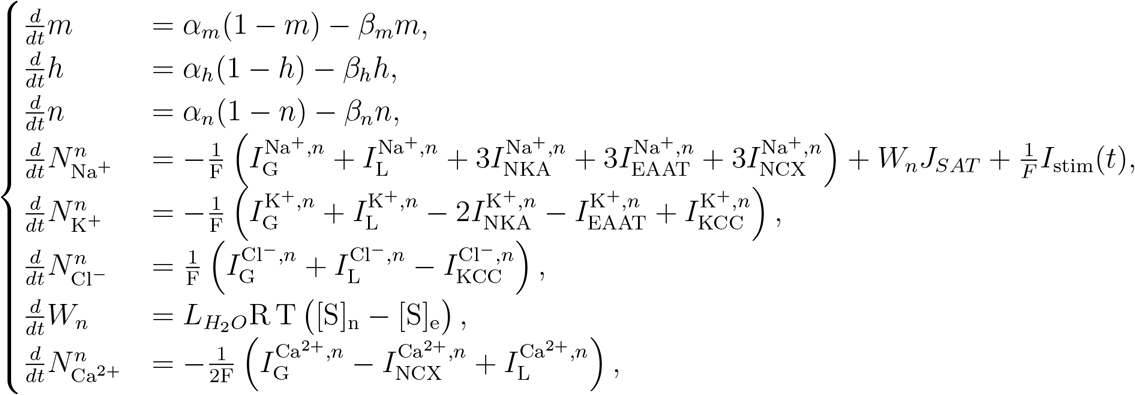

### D.2 Calcium-dependent glutamate cycle

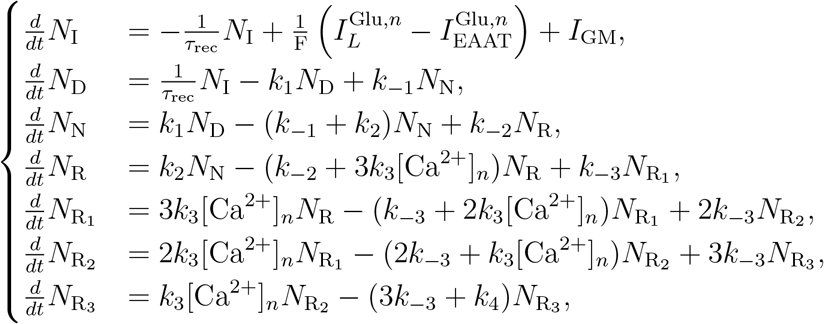

### D.3 Astrocyte

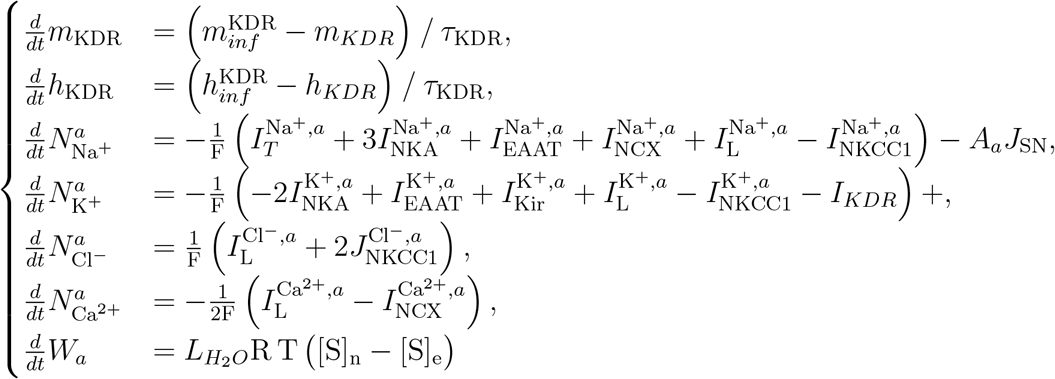

### D.4 Glutamate-glutamine cycle

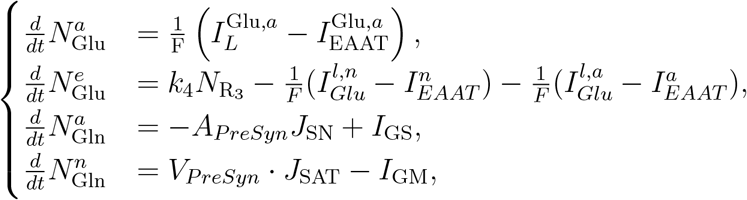

### D.5 Oxygen dynamics

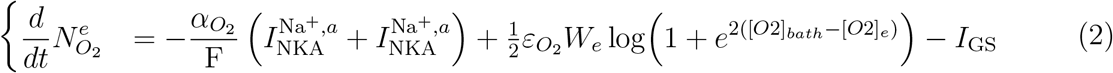

## E Model parameters

**Table 3:** Universal parameters.

| Parameter | Value | Units | Description |
| --- | --- | --- | --- |
| $F$ | 96485.333 | C/mol | Faraday's constant |
| $R$ | 8314.4598 | (C·mV)/(mol·K) | Universal gas constant |
| $T$ | 310 | K | Absolute temperature |
| $Z_{\text{Na}}$ | 1 | – | Valence of Na |
| $Z_{\text{K}}$ | 1 | – | Valence of K |
| $Z_{\text{B}}$ | 1 | – | Valence (unspecified) |
| $Z_{\text{Ca}}$ | 2 | – | Valence of Ca |
| $Z_{\text{Glu}}$ | -1 | – | Valence of Glutamate |
| $Z_{\text{Cl}}$ | -1 | – | Valence of Cl |
| $Z_{\text{A}}$ | -1 | – | Valence (unspecified) |
| $C$ | 20 | pF | Membrane capacitance |
| $R_{H_e, H_a}$ | 2/3 | – | Proton ratio in neurons |
| $L_{\text{H}_2\text{O}, n}$ | $2 \times 10^{-14}$ | m<sup>3</sup>/ms/bar | Neuronal water permeability |
| $L_{\text{H}_2\text{O}, a}$ | $2 \times 10^{-14}$ | m<sup>3</sup>/ms/bar | Astrocytic water permeability |
| $\alpha_{\text{O}_2}$ | 5.3/32 | - | Conversion factor |
| $\varepsilon_{\text{O}_2}$ | $25 \times 10^{-6}$ | ms<sup>-1</sup> | Oxygen diffusion rate |
| $\text{Vol}_{\text{PreSyn}, n}$ | $1 \times 10^{-3}$ | mm<sup>3</sup> | Presynaptic neuron terminal volume |
| $\text{Vol}_{\text{PreSyn}, a}$ | $1 \times 10^{-3}$ | mm<sup>3</sup> | Presynaptic astrocyte terminal volume |
| $\text{Vol}_{\text{Cleft}}$ | $1 \times 10^{-3}$ | mm<sup>3</sup> | Synaptic cleft volume |
| $\text{Radius}_{\text{PreSyn}, n}$ | $(\frac{3 \cdot \text{Vol}_{\text{PreSyn}, n}}{4\pi})^{1/3}$ | mm | Radius of presynaptic neuronal terminal |
| $\text{Radius}_{\text{PreSyn}, a}$ | $(\frac{3 \cdot \text{Vol}_{\text{PreSyn}, a}}{4\pi})^{1/3}$ | mm | Radius of presynaptic astrocytic terminal |
| $\text{Area}_{\text{PreSyn}, n}$ | $4\pi \cdot \text{Radius}_{\text{PreSyn}, n}^2$ | mm<sup>2</sup> | Area of presynaptic neuronal terminal |
| $\text{Area}_{\text{PreSyn}, a}$ | $4\pi \cdot \text{Radius}_{\text{PreSyn}, a}^2$ | mm<sup>2</sup> | Area of presynaptic astrocytic terminal |

**Table 4:** Ion Permeabilities and Conductances.

| Parameter | Value | Units | Description |
| --- | --- | --- | --- |
| **Neuronal Channels** |  |  |  |
| $P_{Na^+}^{t,n}$ | $80 \times 10^{-4}$ | $\text{mm}^3/\text{ms}$ | Transient $\text{Na}^+$ channel permeability |
| $P_{Na^+}^{l,n}$ | $2 \times 10^{-6}$ | $\text{mm}^3/\text{ms}$ | Leak $\text{Na}^+$ permeability |
| $P_{d,K,n}$ | $400 \times 10^{-6}$ | $\text{mm}^3/\text{ms}$ | Delayed rectifier $\text{K}^+$ permeability |
| $P_{K^+}^{l,n}$ | $20 \times 2^{-5}$ | $\text{mm}^3/\text{ms}$ | Leak $\text{K}^+$ permeability |
| $P_{Cl^-}^{g,n}$ | $19.5 \times 10^{-6}$ | $\text{mm}^3/\text{ms}$ | Gated $\text{Cl}^-$ permeability |
| $P_{Cl^-}^{l,n}$ | $2.5 \times 10^{-6}$ | $\text{mm}^3/\text{ms}$ | Leak $\text{Cl}^-$ permeability |
| $P_{Ca^{2+}}^{g,n}$ | $1.6 \times 10^{-5}$ | $\text{mm}^3/\text{ms}$ | Gated $\text{Ca}^{2+}$ permeability |
| $P_{Ca^{2+}}^{l,n}$ | $4 \times 10^{-10}$ | $\text{mm}^3/\text{ms}$ | Leak $\text{Ca}^{2+}$ permeability |
| $P_{Glu}^{l,n}$ | $1 \times 10^{-7}$ | $\text{mm}^3/\text{ms}$ | Leak Glu permeability |
| **Astrocytic Channels** |  |  |  |
| $P_{Ca^{2+}}^{l,a}$ | $4 \times 10^{-11}$ | $\text{mm}^3/\text{ms}$ | Leak $\text{Ca}^{2+}$ permeability |
| $P_{Glu}^{l,a}$ | $1 \times 10^{-10}$ | $\text{mm}^3/\text{ms}$ | Leak Glu permeability |
| $P_{Na^+}^{l,a}$ | $1 \times 10^{-7}$ | $\text{mm}^3/\text{ms}$ | Leak $\text{Na}^+$ permeability |
| $P_{K^+}^{l,a}$ | $70 \times 10^{-6}$ | $\text{mm}^3/\text{ms}$ | Leak $\text{K}^+$ permeability |
| $P_{Cl^-}^{l,a}$ | $1.5 \times 10^{-6}$ | $\text{mm}^3/\text{ms}$ | Leak $\text{Cl}^-$ permeability |
| $g_{\text{KDR}}$ | 3566.4 | — | KDR conductance |
| $P_{\text{KDR}}$ | 40 | — | KDR permeability |

**Table 5:**
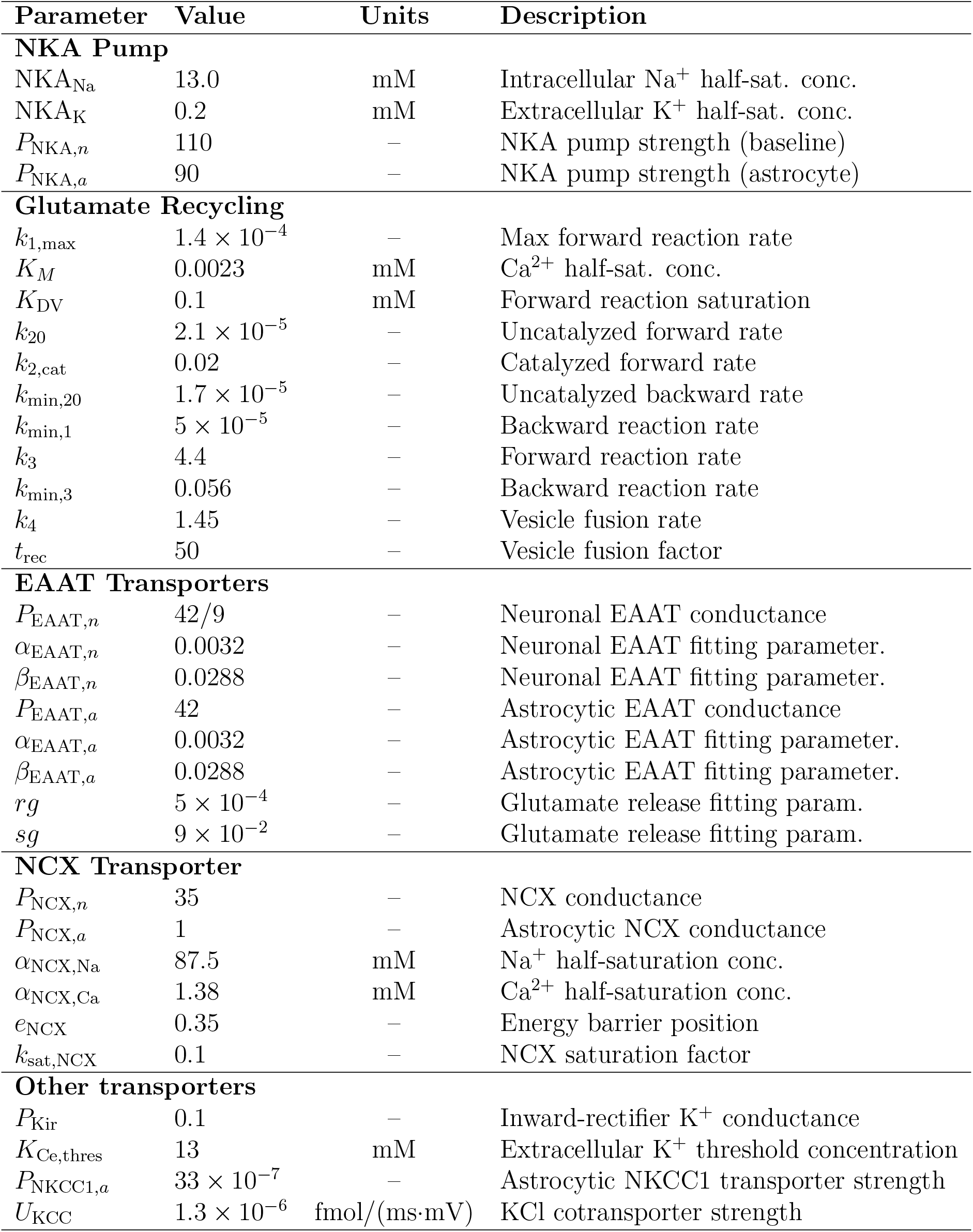
Transporter and Recycling Parameters.

| Parameter | Value | Units | Description |
| --- | --- | --- | --- |
| **NKA Pump** |  |  |  |
| $NKA_{Na}$ | 13.0 | mM | Intracellular $Na^+$ half-sat. conc. |
| $NKA_K$ | 0.2 | mM | Extracellular $K^+$ half-sat. conc. |
| $P_{NKA,n}$ | 110 | – | NKA pump strength (baseline) |
| $P_{NKA,a}$ | 90 | – | NKA pump strength (astrocyte) |
| **Glutamate Recycling** |  |  |  |
| $k_{1,max}$ | $1.4 \times 10^{-4}$ | – | Max forward reaction rate |
| $K_M$ | 0.0023 | mM | $Ca^{2+}$ half-sat. conc. |
| $K_{DV}$ | 0.1 | mM | Forward reaction saturation |
| $k_{20}$ | $2.1 \times 10^{-5}$ | – | Uncatalyzed forward rate |
| $k_{2,cat}$ | 0.02 | – | Catalyzed forward rate |
| $k_{min,20}$ | $1.7 \times 10^{-5}$ | – | Uncatalyzed backward rate |
| $k_{min,1}$ | $5 \times 10^{-5}$ | – | Backward reaction rate |
| $k_3$ | 4.4 | – | Forward reaction rate |
| $k_{min,3}$ | 0.056 | – | Backward reaction rate |
| $k_4$ | 1.45 | – | Vesicle fusion rate |
| $t_{rec}$ | 50 | – | Vesicle fusion factor |
| **EAAT Transporters** |  |  |  |
| $P_{EAAT,n}$ | 42/9 | – | Neuronal EAAT conductance |
| $\alpha_{EAAT,n}$ | 0.0032 | – | Neuronal EAAT fitting parameter. |
| $\beta_{EAAT,n}$ | 0.0288 | – | Neuronal EAAT fitting parameter. |
| $P_{EAAT,a}$ | 42 | – | Astrocytic EAAT conductance |
| $\alpha_{EAAT,a}$ | 0.0032 | – | Astrocytic EAAT fitting parameter. |
| $\beta_{EAAT,a}$ | 0.0288 | – | Astrocytic EAAT fitting parameter. |
| $rg$ | $5 \times 10^{-4}$ | – | Glutamate release fitting param. |
| $sg$ | $9 \times 10^{-2}$ | – | Glutamate release fitting param. |
| **NCX Transporter** |  |  |  |
| $P_{NCX,n}$ | 35 | – | NCX conductance |
| $P_{NCX,a}$ | 1 | – | Astrocytic NCX conductance |
| $\alpha_{NCX,Na}$ | 87.5 | mM | $Na^+$ half-saturation conc. |
| $\alpha_{NCX,Ca}$ | 1.38 | mM | $Ca^{2+}$ half-saturation conc. |
| $e_{NCX}$ | 0.35 | – | Energy barrier position |
| $k_{sat,NCX}$ | 0.1 | – | NCX saturation factor |
| **Other transporters** |  |  |  |
| $P_{Kir}$ | 0.1 | – | Inward-rectifier $K^+$ conductance |
| $K_{Ce,thres}$ | 13 | mM | Extracellular $K^+$ threshold concentration |
| $P_{NKCC1,a}$ | $33 \times 10^{-7}$ | – | Astrocytic NKCC1 transporter strength |
| $U_{KCC}$ | $1.3 \times 10^{-6}$ | fmol/(ms·mV) | KCl cotransporter strength |

**Table 6:** Parameters for Glutamate-Glutamine Cycle and Transporters.

| Parameter | Value | Units | Description |
| --- | --- | --- | --- |
| $\varphi_{\text{GS}}$ | $3 \times 10^{-7}$ | fmol/ms | Glutamine synthetase activity |
| $\varphi_{\text{SN}}$ | 6e-5 | – | SN transporter rate constant |
| $\varphi_{\text{SAT}}$ | $1.7 \times 10^{-4}$ | fmol/mm<sup>3</sup>/ms | SAT transporter rate constant |
| $\varphi_{\text{GM}}$ | $1 \times 10^{-7}$ | fmol/ms | Glutaminase activity |
| $P_{\text{SN}}$ | $1.755 \times 10^{-4}$ | fmol/100$\mu\text{m}^2$·ms | SN transporter permeability |
| $K_{\text{m,GM}}^{\text{Gln}}$ | 0.6 | mM | Michaelis-Menten constant for glutamine |
| $K_{\text{m,GS}}^{\text{Glu}}$ | 2.5 | mM | Michaelis-Menten constant for glutamate |
| $K_{\text{m,GS}}^{\text{NH}_4}$ | 0.2 | mM | Michaelis-Menten constant for NH<sub>4</sub><sup>+</sup> |
| $K_{\text{m,GS}}^{\text{ATP}}$ | 2.3 | mM | Michaelis-Menten constant for ATP |
| $K_{\text{m,SN}}^{\text{Gln}}$ | 1.57 | mM | Michaelis-Menten constant for glutamine |
| $K_{\text{m,SN}}^{\text{Na}}$ | 31 | mM | Michaelis-Menten constant for Na<sup>+</sup> |
| $K_{\text{m,SN}}^{\text{H}}$ | $100 \times 10^{-6}$ | mM | Michaelis-Menten constant for H<sup>+</sup> |
| $K_{\text{m,SAT}}^{\text{Gln}}$ | 0.3 | mM | Michaelis-Menten constant for glutamine |
| $K_{\text{m,SAT}}^{\text{Na}}$ | 10 | mM | Michaelis-Menten constant for Na<sup>+</sup> |
| NH<sub>4,a</sub><sup>+</sup> | 0.15 | mM | Astrocytic ammonium concentration |
| $H_a$ | $6.3096 \times 10^{-5}$ | mM | Astrocytic proton concentration |
| $H_e$ | $6.3069 \times 10^{-5}$ | mM | Extracellular proton concentration |

## Data Availability Statement

The code for all the simulations performed are available at https://github.com/HannahvanSusteren3/GGsyntrans

## Acknowledgements

This study was supported by the funds from the Deutsche Forschungsgemeinschaft (DFG), FOR2795 ‘Synapses under stress’ to CRR (Prof. Dr. Christine R. Rose) (Ro2327/13-2 and 14-2).

## Co-author details

**Hannah van Susteren**, Department of Applied Mathematics, University of Twente, Enschede, the Netherlands.

**Christine R. Rose**, Institute of Neurobiology, Heinrich Heine University, Düsseldorf, Germany.

**Michel J.A.M. van Putten**, Clinical Neurophysiology group, department of Science and Technology, University of Twente, Enschede, the Netherlands and Medisch Spectrum Twente, Enschede, the Netherlands.

**Hil. G.E. Meijer**, Department of Applied Mathematics, University of Twente, Enschede, the Netherlands.

**Author contributions**

**Hannah van Susteren**: Conceptualization, Investigation, Methodology, Software, Validation, Visualization, Writing – Original Draft Preparation, Writing – Review & Editing.

**Christine R. Rose**: Conceptualization, Funding Acquisition, Writing – Review & Editing.

**Michel J.A.M. van Putten**: Conceptualization, Funding Acquisition, Supervision, Writing – Original Draft Preparation, Writing – Review & Editing.

**Hil G.E. Meijer**: Conceptualization, Funding Acquisition, Methodology, Supervision, Writing – Original Draft Preparation, Writing – Review & Editing.

All authors contributed to the article and approved the submitted version.

